# Distinct metabolic states of yeast and pseudohyphae differentially prime innate immune responsiveness

**DOI:** 10.64898/2026.10.08.757614

**Authors:** Hyuga Kato, Monali NandyMazumdar, Zahra Rahmani, Teagan Kukhta, Claire Wolford, Sahishnu Vallabhajoysul, Masaru Miyagi, Susumu Kajiwara, Mei Zhang

**Affiliations:** Department of Biomedical Engineering, Case Western Reserve University School of Medicine, Cleveland, OH, USA; Case Comprehensive Cancer Center, Cleveland, OH, USA; Department of Life Science and Technology, Tokyo Institute of Science, Tokyo, Japan; Department of Computer and Data Sciences, Case Western Reserve University School of Engineering, Cleveland, OH, USA; Department of Genetics and Genome Sciences, BSTP program, Case Western Reserve University School of Medicine, Cleveland, OH 44106, USA; Department of Pharmacology, Case Western Reserve University School of Medicine, Cleveland, OH, USA

**Keywords:** *Malassezia furfur*, fungal morphogenesis, fungal metabolic states, tryptophan aldehyde metabolism, metabolomics, host–microbe interactions, innate immunity, immunometabolism

## Abstract

Pseudohyphal growth in skin-associated fungi is often considered a morphological response to environmental stress, but whether it represents a coordinated metabolic state that influences host immunity remains unclear. Using *Malassezia furfur*, we combined synchronized morphogenesis, temporal metabolomics, stable-isotope tracing, targeted metabolite measurements, transcriptional profiling, and functional immune assays to compare yeast and pseudohyphal states. Yeast cells maintained an active tryptophan-aldehyde metabolic program that partitioned carbon between oxidative and reductive branches associated with redox balance and methylation potential. Pseudohyphal cells attenuated this program and instead underwent sequential metabolic remodeling, characterized by early changes in one-carbon and amino acid metabolism followed by sulfur-pathway remodeling and increased glutathione biosynthesis. These distinct fungal states generated extracellular environments that altered THP-1 innate immune responsiveness without directly inducing canonical macrophage polarization. Yeast-conditioned cells responded preferentially to tissue-repair signals, whereas pseudohypha-conditioned cells displayed enhanced responses to inflammatory stimuli. Individual metabolites did not reproduce these effects, suggesting that immune conditioning depends on the integrated extracellular milieu. Together, these findings identify pseudohyphal growth as an organized adaptive metabolic state and establish a framework linking fungal morphogenetic potential to context-dependent host immune responses.

**Significance Statement:** Fungal morphology is commonly interpreted as a marker of development or virulence, yet pseudohyphal growth in skin-associated fungi may instead reflect an adaptive physiological state. Using *Malassezia furfur*, we show that yeast and pseudohyphal cells engage distinct metabolic programs that generate different extracellular environments and differentially condition innate immune responsiveness. Rather than directly polarizing macrophages, yeast-associated environments favor responsiveness to tissue-repair cues, whereas pseudohypha-associated environments increase responsiveness to inflammatory stimuli. These effects are not reproduced by individual metabolites, indicating that they arise from the integrated fungal extracellular milieu. Our findings suggest that fungal physiological state and morphogenetic potential, rather than fungal abundance alone, influence host immune outcomes and may help explain the variable contribution of *Malassezia* to chronic inflammatory skin disease.

## Introduction

*Malassezia* species are prominent members of the human skin mycobiota and have been associated with pityriasis versicolor, seborrheic dermatitis, atopic dermatitis, and folliculitis (1–4). However, these fungi also colonize healthy skin, and disease-associated strains can be present at nonlesional sites (5,6). Their presence or abundance alone therefore does not adequately explain the variability of host responses. A more informative framework may require consideration of fungal physiological state together with the surrounding host environment.

Fungal morphology provides a visible indication of underlying physiological organization. Yeast, pseudohyphal, and true hyphal forms represent distinct morphogenetic programs rather than successive points along a single developmental continuum (7–9). Pseudohyphae consist of elongated, attached cells with constrictions at their septal junctions, whereas true hyphae display sustained polarized growth and specialized filamentous architecture (7–9). Although true hyphal differentiation has been extensively investigated in relation to development and virulence (10, 11), pseudohyphal growth frequently accompanies environmentally responsive metabolic adaptation (12, 13). Its reversibility and context dependence suggest that it should be evaluated as a distinct physiological state rather than simply as an incomplete hyphal phenotype.

The capacity for pseudohyphal growth also varies among fungal species and strains, indicating that morphological responses are not dictated solely by environmental exposure. Among skin-associated fungi, *Malassezia furfur*, *M. globosa*, and *M. sympodialis* can undergo pseudohyphal transitions, whereas *M. slooffiae* and *M. japonica* do not under comparable conditions (14–17). Related variation is also observed among *Candida* species: *C. auris* can produce stress-associated pseudohyphae, whereas *C. albicans* is also capable of true hyphal differentiation (18–20). Such differences raise the possibility that intrinsic morphogenetic potential contributes to variation in fungal physiology and host interactions.

Cellular metabolism provides a plausible link between fungal morphology and host response because adaptive changes in biosynthesis, redox balance, nutrient utilization, and antioxidant defense can reshape the extracellular milieu (19–23). Although individual fungal metabolites have been implicated in host-microbe communication, it remains unclear whether pseudohyphal growth involves coordinated remodeling across interconnected metabolic pathways and whether the resulting extracellular environment alters innate immune responsiveness.

*M. furfur* offers an experimentally tractable system for examining these questions because selected strains can be maintained in yeast and pseudohyphal states under controlled conditions (12, 14–17). Establishing both morphotypes from the same starting inoculum and subsequently transferring them into a common nutrient environment permits comparison of state-associated metabolic trajectories within a shared genetic and experimental background. We used this platform to integrate temporal metabolomics, stable-isotope tracing, targeted metabolite measurements, transcriptional analysis, and functional immune assays.

We asked whether yeast and pseudohyphal cells occupy distinct metabolic states, how those states develop over time, and whether their extracellular environments alter subsequent innate immune responses. We further considered whether variation in morphogenetic propensity and the stability of pseudohyphal growth could help explain heterogeneous *Malassezia*-associated immune phenotypes. This approach shifts the focus from fungal detection or burden alone to the functional state through which the fungus interacts with its host.

## Results

### Temporal metabolic trajectory analysis reveals distinct metabolic states associated with pseudohyphal growth

To determine whether pseudohyphal growth in *M. furfur* is associated with a distinct metabolic state, we established a synchronized morphogenesis platform that generated yeast and pseudohyphal populations from the same starting inoculum (Fig. 1 A-C). After the two morphotypes were established, cultures were normalized to equivalent cell densities, transferred into a common nutrient environment, and sampled over time to characterize their metabolic trajectories (Fig. 1D). This design enabled comparison of morphotype-associated physiology within a shared genetic background and under identical subsequent culture conditions.

**Figure. 1.**
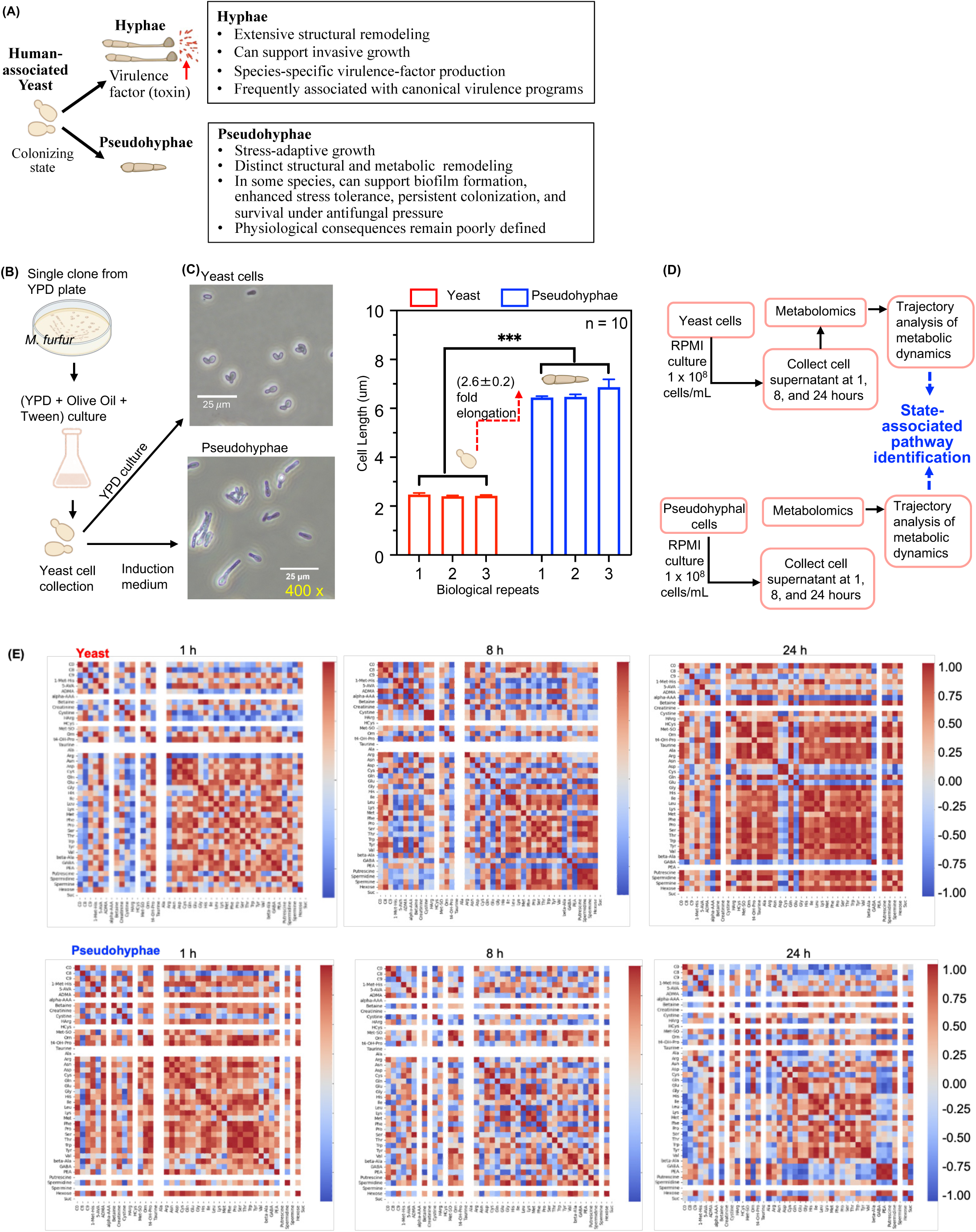
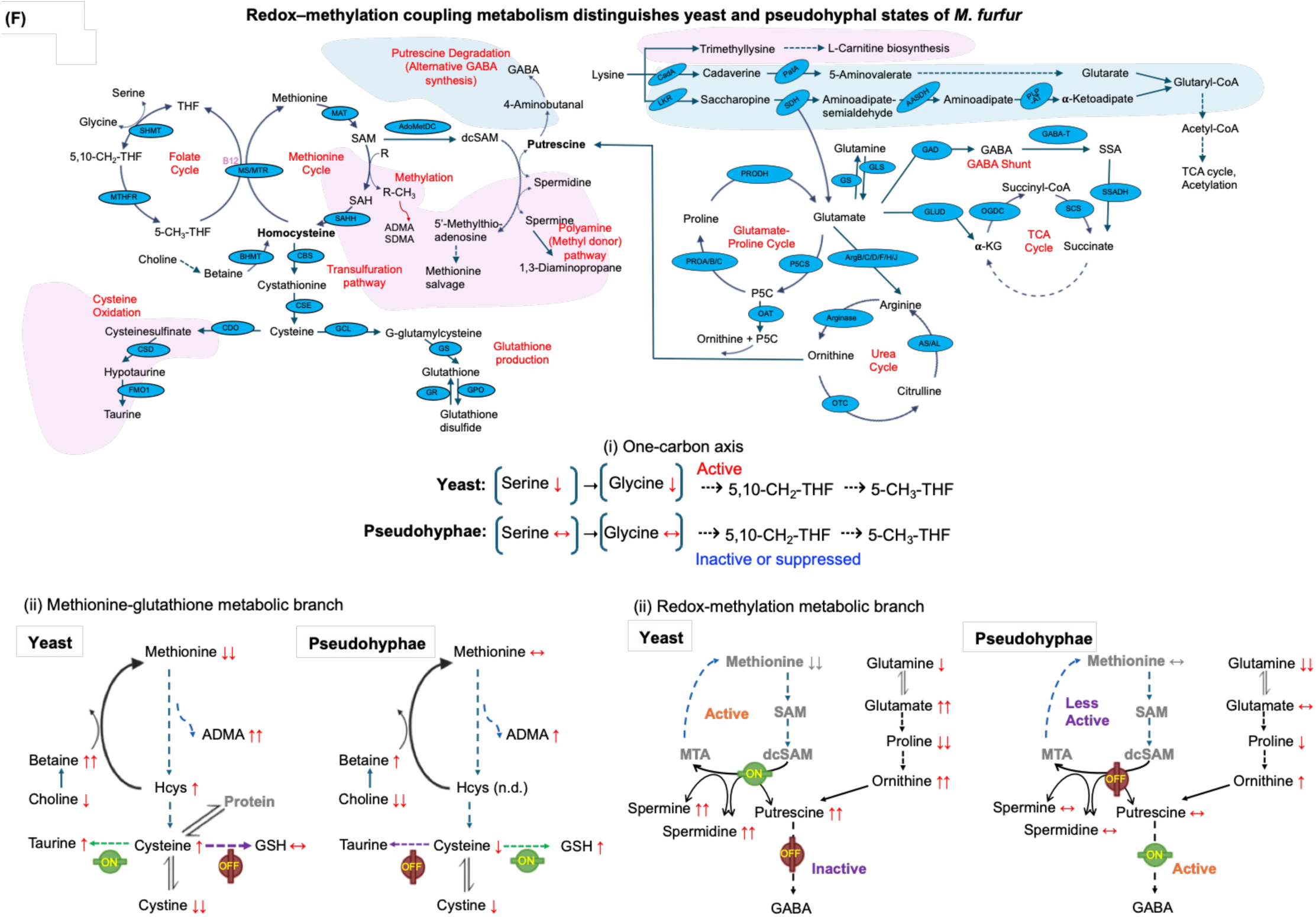
Morphogenesis establishes distinct fungal metabolic states through temporal metabolomic trajectory analysis. **(A)** Conceptual comparison of true hyphal and pseudohyphal morphogenesis in human-associated fungi, including species of *Candida* and *Malassezia*. **(B)** Schematic of the synchronized yeast and pseudohyphal cell culture platform. *M. furfur* CBS1878 cells were cultured under defined conditions to maintain the yeast state or induce pseudohyphal differentiation. **(C)** Microscopic morphological characterization of yeast and pseudohyphal states. Representative bright-field images show unicellular yeast cells and elongated pseudohyphal structures following induction (scale bars, 25 μm). Cell length was quantified to compare the two morphotypes. **(D)** Experimental design for temporal metabolomic analysis. Established yeast and pseudohyphal cells were harvested, normalized to equivalent cell densities, transferred into serum-free RPMI medium, and sampled at defined time points with four biological replicates at each time points for extracellular metabolomic profiling. N = 4 **(E)** Unsupervised correlation-based trajectory analysis of temporal metabolomic profiles. Distinct temporal metabolic trajectories were identified for yeast and pseudohyphal cultures based on coordinated metabolite correlation patterns. An overview of the metabolite classes detected by temporal metabolomic profiling is provided in Supplementary Fig. S1, and the complete quantitative metabolomic dataset across morphotypes and time points is provided in Supplementary Table S1. **(F)** Integrated pathway analysis of morphotype-associated metabolic organization. Coordinated remodeling was observed across interconnected pathways including one-carbon metabolism, methionine cycling, transmethylation reactions, sulfur amino acid metabolism, glutathione metabolism, and central carbon metabolism (Supplementary Fig S2 A-C). Independent targeted LC–MS/MS quantification of formate and reduced and oxidized glutathione (GSH and GSSG) is presented in Supplementary Fig. S2 D and E. Panels. Panels i–iii provide a summary model illustrating morphogenesis-associated fungal metabolic states. The concentration trajectories of all metabolites shown in panels i-iii are provided in Supplementary Fig. S2. Following transfer into an identical nutrient environment, synchronized yeast and pseudohyphal cells exhibited distinct temporal metabolic organization.

Unsupervised, correlation-based analysis identified distinct temporal patterns in yeast and pseudohyphal cultures, with coordinated differences spanning multiple interconnected metabolic networks (Fig. 1E, Supplementary Fig. S1, and Supplementary Table S1). Representative metabolite trajectories differed in serine and glycine utilization, methionine cycling, polyamine turnover, cysteine metabolism, and glutathione accumulation (Supplementary Fig. S2 A-C). Independent targeted LC-MS/MS measurements of formate, reduced glutathione (GSH), and oxidized glutathione (GSSG) further supported differences in one-carbon metabolism and glutathione-associated redox responses (Supplementary Fig. S2 D and E).

Pathway-level integration localized these coordinated changes to one-carbon metabolism, methionine cycling, transmethylation, polyamine metabolism, sulfur amino acid metabolism, glutathione metabolism, and central carbon metabolism (Fig. 1F). Thus, yeast and pseudohyphal cultures occupy distinguishable metabolic states defined by network-level remodeling.

### Pseudohyphal metabolic remodeling develops in a temporally ordered sequence

We next examined whether metabolic differences between the two morphotypes appeared simultaneously or emerged in a defined sequence using metabolomic trajectory analysis. By 8 h, metabolite ratios associated with one-carbon metabolism, methyl-donor availability, amino acid utilization, and oxidative metabolism had already diverged (Fig. 2A). These included betaine/choline, serine/glycine, methionine sulfoxide/methionine, glutamate/glutamine, tyrosine/phenylalanine, aminoadipate/lysine, hydroxyproline/proline, and asymmetric dimethylarginine/arginine, indicating early differences across interconnected metabolic processes.

**Fig. 2.**
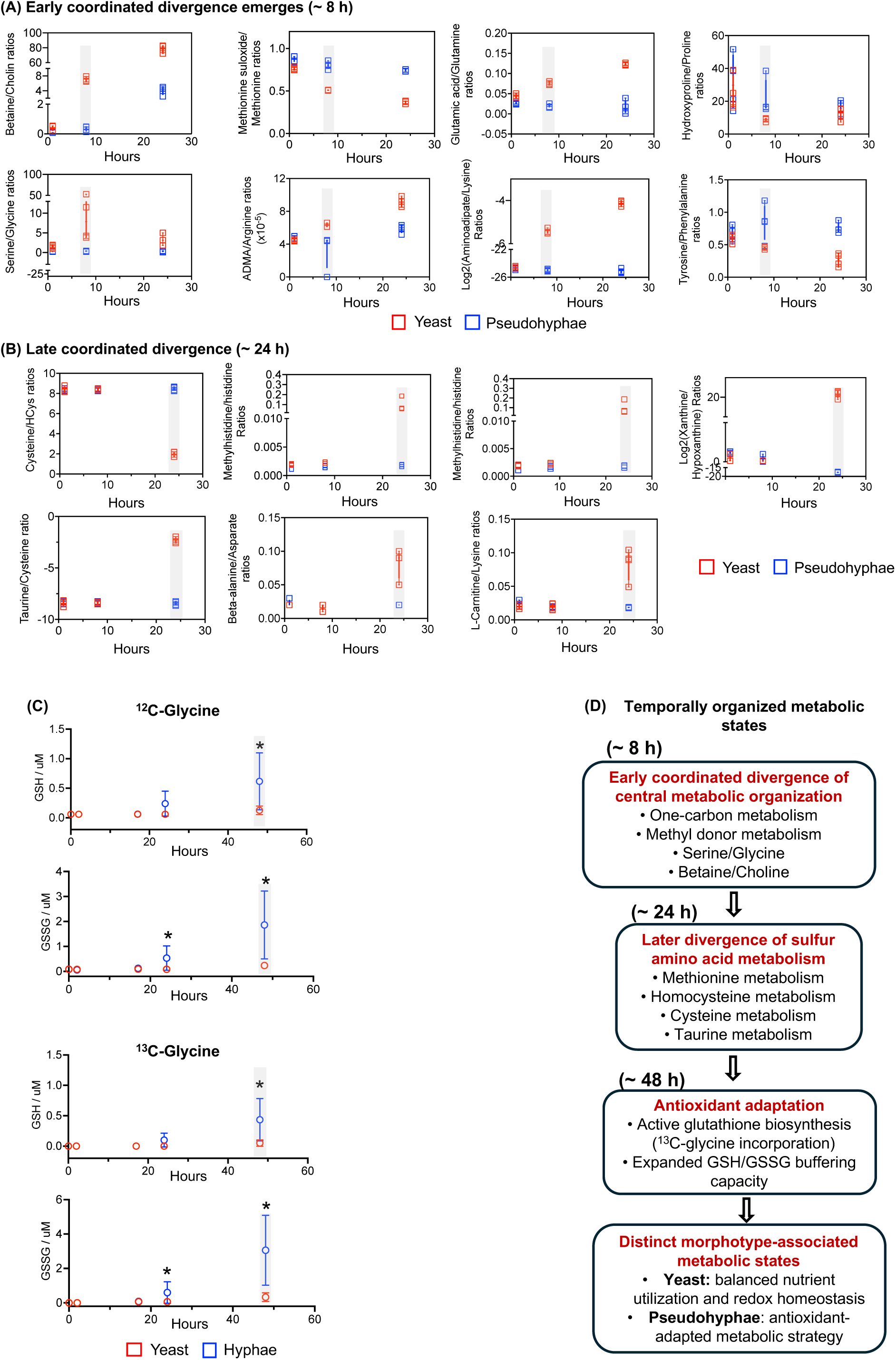
Sequential temporal remodeling of interconnected metabolic pathways distinguishes nutrient utilization by established fungal metabolic states. **(A)** Ratio-based temporal metabolomic analysis of conditioned media. Representative metabolite ratios associated with one-carbon metabolism, methyl donor metabolism, amino acid utilization, and oxidative metabolism exhibit the earliest divergence between established east and pseudohyphal cultures following transfer into RPMI. Gray shading indicates the time point (∼8 h) at which morphotype-associated metabolic trajectories become distinguishable. **(B)** Temporal remodeling of sulfur amino acid-associated metabolic pathways. Representative metabolite ratios associated with cysteine–homocysteine metabolism, taurine biosynthesis, purine metabolism, β-alanine metabolism, histidine methylation, and L-carnitine metabolism diverge during later stages of culture. Gray shading indicates the onset of coordinated pathway divergence (∼24 h). **(C)** Functional analysis of glutathione metabolism. Targeted LC–MS/MS quantification following supplementation with naturally abundant (¹²C) glycine shows accumulation of reduced glutathione (GSH) and oxidized glutathione (GSSG) in yeast and pseudohyphal cultures at 24–48 h. Parallel stable isotope tracing using uniformly labeled (¹³C) glycine demonstrates incorporation of extracellular glycine into newly synthesized GSH and GSSG. Gray shading indicates the later emergence of glutathione metabolic remodeling between yeast and pseudohyphal cultures. **(D)** Summary model illustrating the temporal organization of pathway-specific metabolic trajectories during nutrient utilization by established yeast and pseudohyphal fungal states. Early divergence of central metabolic organization is followed by remodeling of sulfur amino acid metabolism and subsequent expansion of glutathione metabolic remodeling, resulting in distinct morphotype-associated metabolic trajectories.

Additional divergence became evident at approximately 24 h, including changes associated with cysteine-homocysteine metabolism, taurine biosynthesis, purine metabolism, β-alanine metabolism, histidine methylation, and L-carnitine metabolism (Fig. 2B). The delayed emergence of these differences indicates that pseudohyphal metabolic remodeling proceeds from early changes in central metabolic organization to later remodeling of sulfur-associated and related pathways.

Because glutathione connects sulfur metabolism to antioxidant defense, we next assessed whether late pathway remodeling was accompanied by increased glutathione production. Targeted LC-MS/MS demonstrated greater accumulation of both GSH and GSSG in pseudohyphal cultures after 24 h (Fig. 2C). Increased incorporation of ¹³C-glycine into both glutathione species supported active de novo synthesis, identifying enhanced glutathione-associated metabolism as a later feature of the pseudohyphal state.

These findings establish a temporal sequence in which early divergence in central metabolic pathways precedes sulfur-associated remodeling and increased glutathione biosynthesis in pseudohyphal cultures (Fig. 2D).

### Tryptophan-aldehyde metabolism distinguishes yeast state and is attenuated in pseudohyphal cells

To identify metabolic features that distinguished the two morphotypes throughout the time course, we examined ratios that remained consistently different between conditions. The ratio of indole-3-acetic acid (IAA) to tryptophan showed sustained separation from our metabolomic profiling study (Fig. 3A), implicating tryptophan-derived aldehyde metabolism, a pathway with oxidative and reductive branches connected to NAD⁺/NADH balance (Fig. 3B). Oxidation of indole-3-acetaldehyde (IAAld) to IAA reduces NAD⁺ to NADH, whereas reduction of IAAld to tryptophol (TOL) consumes NADH and regenerates NAD⁺. Reduction of indole-3-pyruvate to indole-3-lactic acid (ILA) likewise consumes reducing equivalents. The distribution of tryptophan-derived carbon among these products therefore provides insight into the organization of redox-linked metabolic pathways.

**Figure 3.**
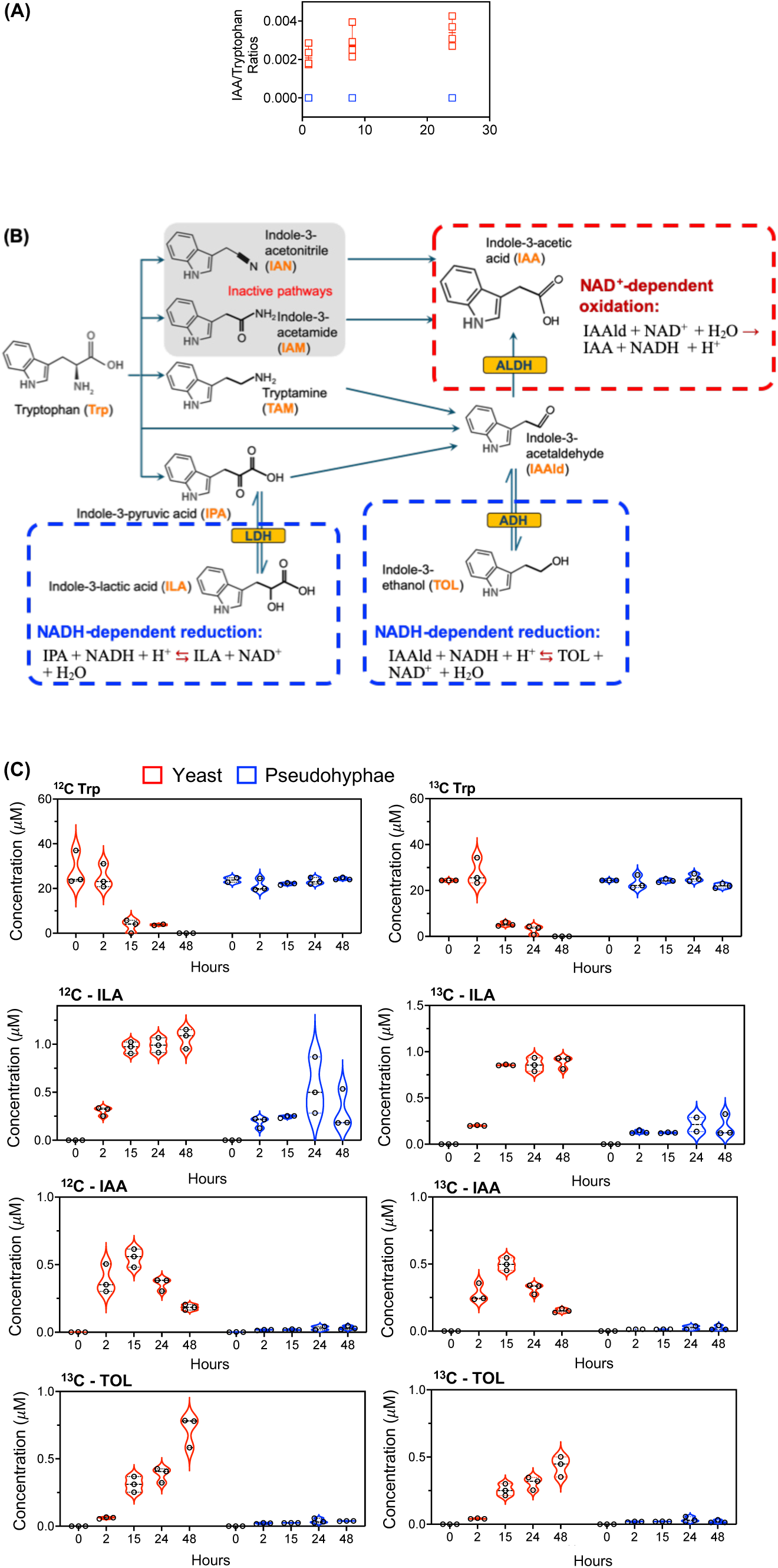
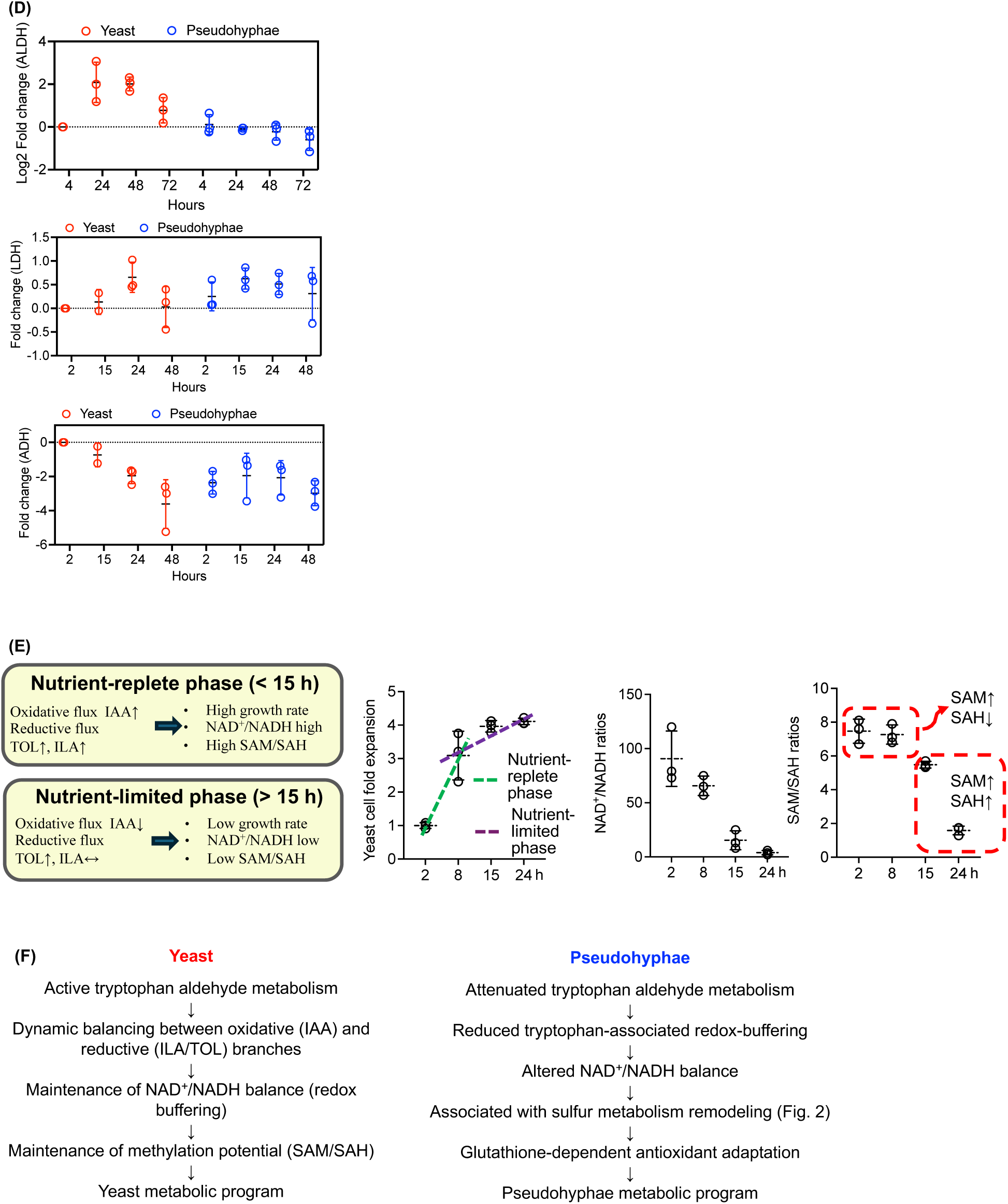
Coordinated partitioning of tryptophan aldehyde metabolism distinguishes the yeast-associated fungal metabolic state. **(A)** Temporal metabolite ratio analysis identifies the indole-3-acetic acid (IAA)/tryptophan ratio as a persistent discriminator between established yeast and pseudohyphal morphotypes cultures throughout nutrient utilization, in contrast to the progressive metabolic remodeling observed across other metabolic pathways (Fig. 2). **(B)** Schematic of tryptophan aldehyde metabolism in *M. furfur*. Tryptophan is metabolized through the indole-pyruvate/indole-3-acetaldehyde pathway, whereas no detectable endogenous or ¹³C-labeled metabolites corresponding to the indole-3-acetamide or indole-3-acetonitrile branches were observed under the experimental conditions (Supplementary Fig. S3). Indole-3-acetaldehyde (IAAld) is converted to indole-3-acetic acid (IAA) through the oxidative branch or to indole-3-lactic acid (ILA) and tryptophol (TOL) through the reductive branches. **(C)** Stable isotope tracing using uniformly labeled ¹³C₁₁-tryptophan. Yeast cultures exhibit robust incorporation of ¹³C into IAA, ILA, and TOL, whereas pseudohyphal cultures display markedly reduced labeling of all three metabolites, indicating attenuation of coordinated tryptophan aldehyde metabolism during pseudohyphal growth. **(D)** Temporal expression of enzymes associated with tryptophan aldehyde metabolism. ALDH transcript abundance closely parallels temporal IAA production, whereas temporal expression of ADH and LDH does not consistently parallel accumulation of TOL and ILA, respectively. **(E)** Temporal organization of the yeast-associated tryptophan aldehyde metabolic program. During the early phase of nutrient utilization (<15 h), coordinated oxidative (IAA) and reductive (ILA/TOL) metabolism is associated with rapid growth, higher NAD⁺/NADH ratios, and elevated methylation potential (SAM/SAH). During later stages of nutrient utilization (>15 h), oxidative metabolism declines relative to reductive metabolism and is accompanied by reduced NAD⁺/NADH ratios and lower methylation potential. **(F)** Summary model illustrating morphotype-associated organization of tryptophan aldehyde metabolism. Yeast cells maintain coordinated partitioning of oxidative and reductive metabolism throughout nutrient utilization, whereas pseudohyphal cells exhibit broad attenuation of tryptophan aldehyde metabolism together with establishment of an alternative metabolic organization characterized by sulfur metabolic remodeling and glutathione-associated adaptation.

Stable-isotope tracing with uniformly labeled ¹³C₁₁-tryptophan showed incorporation of labeled carbon into both oxidative (IAA) and reductive (ILA and TOL) products in yeast cultures, whereas labeling of all three products was reduced in pseudohyphal cultures (Fig. 3C). Measurements of preexisting ¹²C and newly synthesized ¹³C metabolites supported active tryptophan-aldehyde metabolism in yeast cells and coordinated attenuation of its oxidative and reductive branches in pseudohyphal cells.

Transcriptional measurements were consistent with this metabolic organization. Aldehyde dehydrogenase (ALDH) transcript abundance followed the temporal pattern of IAA production, whereas increases in ADH and LDH preceded accumulation of TOL and ILA, respectively (Fig. 3D). Together, these observations link expression of aldehyde-processing enzymes with morphotype-specific differences in tryptophan-derived metabolite production.

During the nutrient-replete phase (<15 h), simultaneous oxidative and reductive product formation coincided with rapid growth and higher S-adenosylmethionine/S-adenosylhomocysteine (SAM/SAH) ratios, consistent with increased methylation potential (Fig. 3E). As nutrient availability declined (>15 h), oxidative product formation decreased while reductive product formation increased, accompanied by changes in NAD⁺/NADH balance. These patterns indicate that tryptophan-aldehyde metabolism in yeast cells is dynamically regulated during adaptation to changing nutrient conditions.

Therefore, active partitioning of tryptophan-derived carbon between oxidative and reductive products characterizes the yeast-associated state, whereas pseudohyphal cells attenuate this program while increasing sulfur-associated and glutathione-linked metabolism (Figs. 2 and 3F).

### Pseudohyphal cells sustain mitochondrial, genome-maintenance, and antioxidant-associated programs

Because reduced tryptophan-aldehyde metabolism may alter redox requirements, we examined transcriptional programs associated with adaptation to the pseudohyphal state. Pseudohyphal cultures showed sustained expression of genes related to mitochondrial function *(IDH2*, *NDUFS4*, and *NDE1*), genome maintenance (*DNA helicase*, *HUS5*, and *INI1*), and glutathione biosynthesis (*GCL* and *GCS*) (Fig. 4).

**Figure 4.**
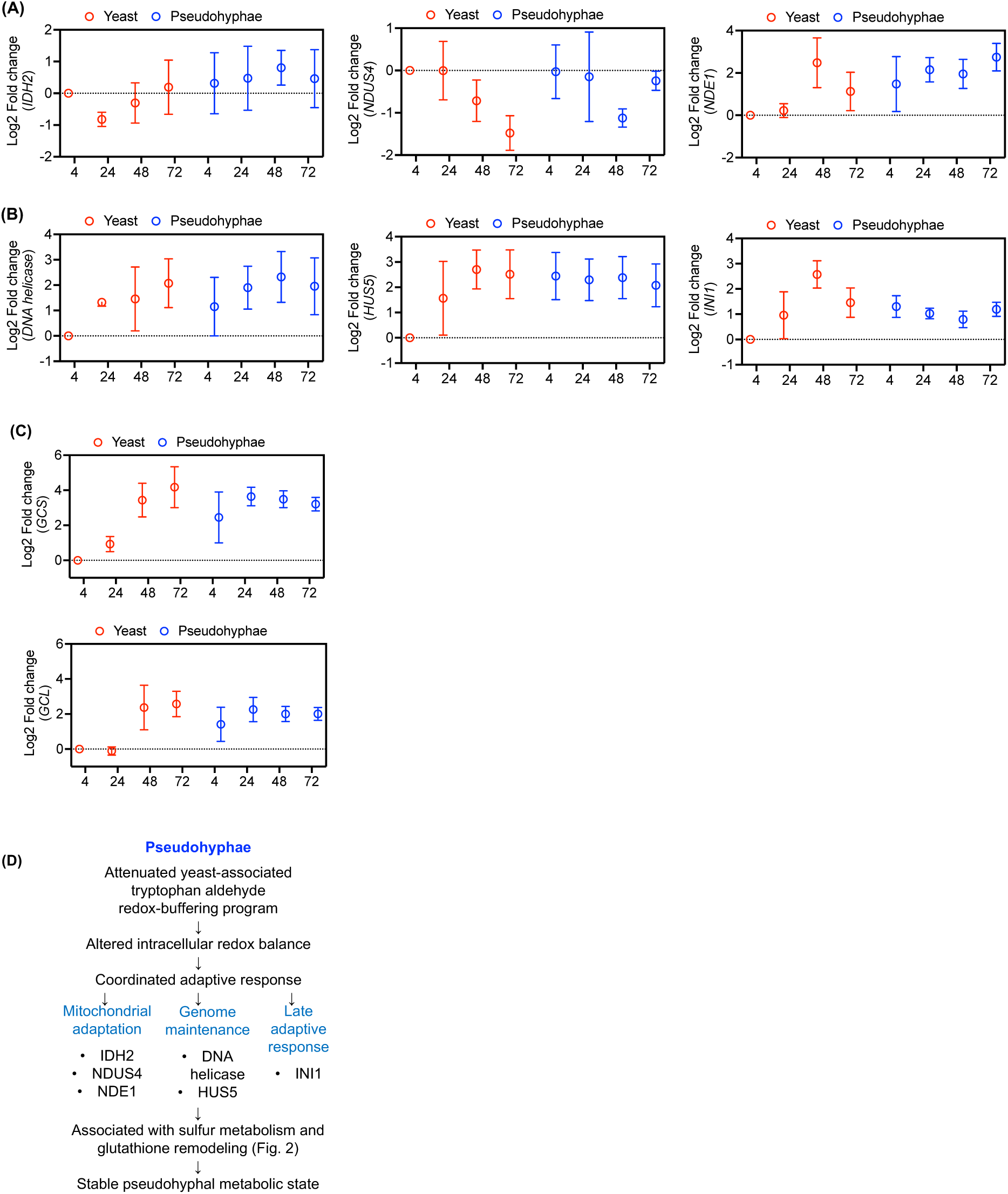
Representative intracellular adaptive responses accompany stabilization of the pseudohyphal metabolic state. **(A)** Temporal expression of representative genes associated with mitochondrial metabolism and respiratory adaptation. Representative genes including IDH2, NDUFS4, and NDE1 exhibit distinct temporal expression trajectories between established yeast and pseudohyphal cultures during nutrient utilization. Pseudohyphal cultures maintain more sustained expression of these representative genes, whereas yeast cultures display transient or declining expression patterns. **(B)** Temporal expression of representative genes associated with genome maintenance and intracellular adaptation. DNA helicase, HUS5, and INI1 exhibit distinct morphotype-dependent temporal expression trajectories during nutrient utilization, with pseudohyphal cultures maintaining more sustained expression than yeast cultures. **(C)** Temporal expression of glutathione biosynthetic genes. Expression of glutamate-cysteine ligase (GCL) and glutathione synthetase (GCS) was quantified throughout nutrient utilization. Pseudohyphal cultures exhibit elevated expression at the initial time points following transfer into RPMI and maintain relatively stable expression thereafter, whereas yeast cultures progressively induce both genes during later stages of nutrient utilization. **(D)** Summary model illustrating representative intracellular adaptive responses accompanying stabilization of the pseudohyphal metabolic state. Attenuation of tryptophan aldehyde metabolism is accompanied by distinct temporal expression of representative genes associated with mitochondrial metabolism, genome maintenance, intracellular adaptation, and glutathione biosynthesis.

*IDH2*, *NDUFS4*, and *NDE1* exhibited transient or declining expression in yeast cultures but remained more consistently expressed in pseudohyphal cultures (Fig. 4A). Similarly, *DNA helicase*, *HUS5*, and *INI1* showed greater temporal variation in yeast cells and comparatively stable expression in pseudohyphal cells (Fig. 4B). The glutathione-biosynthetic genes encoding glutamate-cysteine ligase (*GCL*) and glutathione synthetase (*GCS*) were elevated early in pseudohyphal cultures and remained expressed throughout nutrient utilization, whereas their induction occurred later in yeast cultures (Fig. 4C). These transcriptional patterns preceded the later expansion of GSH and GSSG pools and the increased incorporation of ¹³C-glycine into glutathione.

The combined transcriptional and metabolic profiles associate pseudohyphal growth with sustained mitochondrial adaptation, genome-maintenance responses, and establishment of glutathione-biosynthetic capacity. These features are consistent with an alternative adaptive state accompanying attenuation of tryptophan-aldehyde metabolism.

### Morphotype-associated extracellular environments condition distinct innate immune responses

To determine whether fungal metabolic state influences host cells, we exposed THP-1 monocytes to extracellular conditioned media collected after yeast- and pseudohypha-associated metabolic profiles had been established. Continuous LiCellMo monitoring showed that pseudohypha-conditioned medium induced greater glycolytic activity than yeast-conditioned medium (Fig. 5A). However, neither preparation produced the canonical macrophage morphology observed in cytokine-defined reference populations (Supplementary Fig. S4).

**Figure 5.**
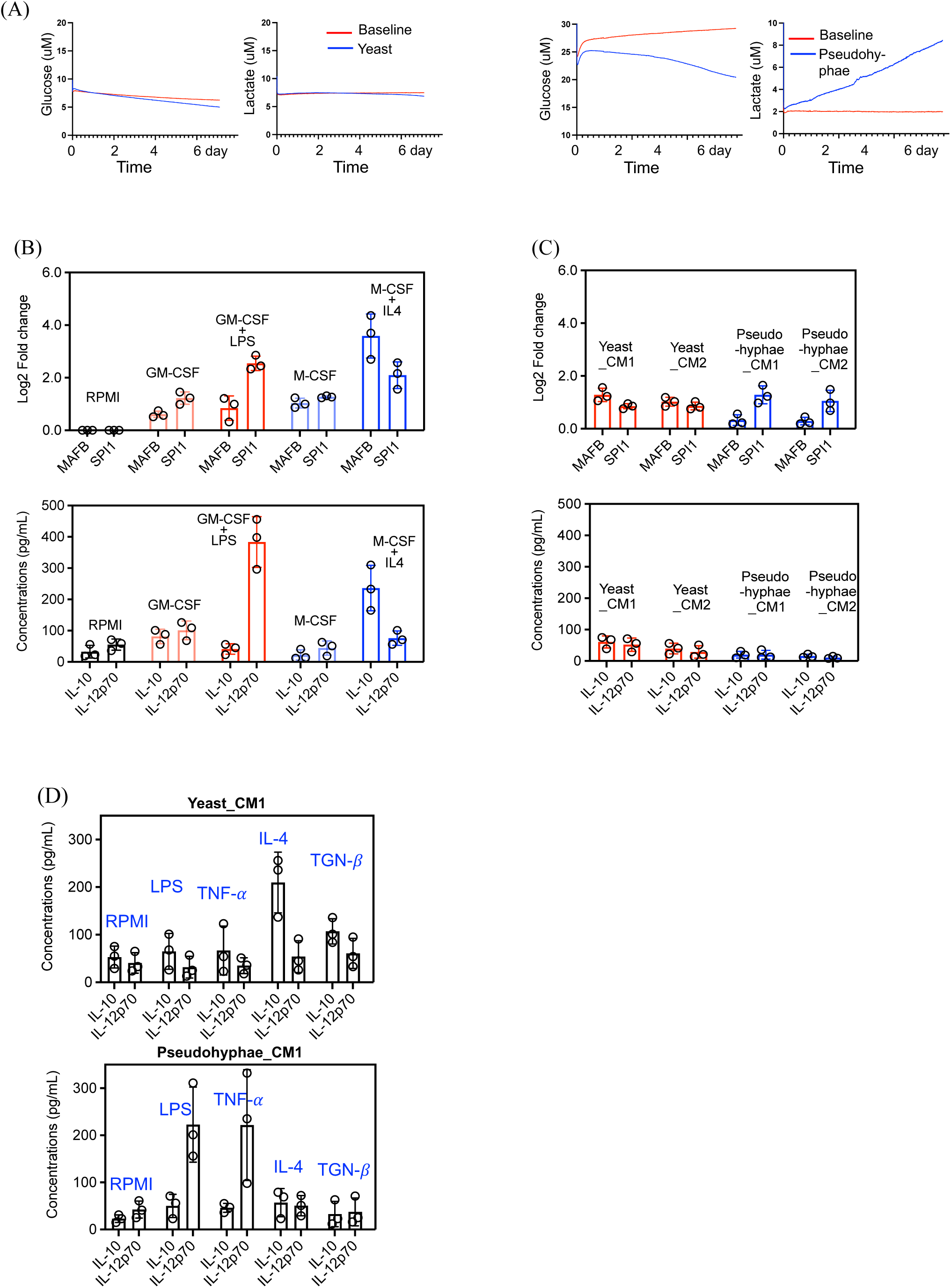
Fungal metabolic states generate extracellular metabolic environments that establish distinct host innate immune response states. **(A)** Real-time metabolic monitoring of THP-1 cells using the LiCellMo platform. Glycolytic activity was continuously quantified by extracellular glucose consumption and lactate production throughout the culture period. THP-1 cells exposed to pseudohyphal extracellular metabolic environments exhibited greater glycolytic activity than cells cultured with yeast-derived extracellular metabolic environments. **(B)** Validation of THP-1 macrophage maturation and polarization reference states. THP-1 cells were cultured under defined cytokine conditions including GM-CSF and M-CSF for macrophage maturation, GM-CSF plus LPS for inflammatory activation, and M-CSF plus IL-4 for tissue-repair activation. Upper panel: RT-qPCR analysis of the macrophage lineage-associated transcription factors MAFB and SPI1 (PU.1). Lower panel: ELISA quantification of IL-12p70 and IL-10. Flow cytometric characterization of these reference populations is presented in Supplementary Fig. S5. **(C)** THP-1 responses following exposure to extracellular metabolic environments generated by established yeast and pseudohyphal fungal metabolic states. Extracellular metabolic environments were used either directly (CM1) or diluted 1:1 with fresh RPMI (CM2). Upper panel: RT-qPCR analysis of MAFB and SPI1 (PU.1). Lower panel: ELISA quantification of IL-10 and IL-12p70. Flow cytometric analyses of CD14, CD80, CD86, CD206, and CD163 are presented in Supplementary Fig. S6. **(D)** Secondary stimulation assay evaluating host innate immune responsiveness following conditioning by fungal extracellular metabolic environments. THP-1 cells were challenged with the pro-inflammatory stimuli LPS or TNF-α or the tissue-repair stimuli IL-4 or TGF-β following conditioning with yeast- or pseudohypha-derived extracellular metabolic environments. Yeast-conditioned cells preferentially responded to tissue-repair stimulation through IL-10 production, whereas pseudohyphal-conditioned cells preferentially responded to pro-inflammatory stimulation through IL-12p70 production.

Reference treatments established inflammatory (GM-CSF/LPS) and tissue-repair-associated (M-CSF/IL-4) macrophage states (Fig. 5B, Supplementary Fig. S5). In comparison, yeast- and pseudohypha-conditioned media induced lineage-associated changes, including reduced CD14 and increased MAFB and SPI1, but did not reproduce either canonical polarization profile or induce basal IL-10 or IL-12p70 production (Fig. 5C, Supplementary Fig. S6). Representative metabolites, including IAA, ILA, TOL, and glutathione, also failed to reproduce the effects of the complete conditioned media (Supplementary, Fig. S7). These results indicate that fungal extracellular environments alter metabolic and lineage-associated properties without directly imposing a defined inflammatory or tissue-repair state.

The functional consequences of this conditioning became apparent after secondary stimulation. Yeast-conditioned cells produced IL-10 in response to IL-4 but showed limited responses to LPS or TNF-α. In contrast, pseudohypha-conditioned cells produced IL-12p70 after stimulation with LPS or TNF-α but showed minimal IL-10 induction in response to IL-4 or TGF-β (Fig. 5D). Thus, the two fungal extracellular environments show distinct response states that favor subsequent responsiveness to tissue-repair-associated or inflammatory signals, respectively.

### Independent studies are consistent with a fungal state-dependent model of host immune responsiveness

To assess whether this framework could inform interpretation of independent studies, we reexamined published datasets describing host responses to *Malassezia* under different morphological or culture conditions. In a human skin explant dataset, *M. sympodialis* maintained a yeast morphology in the absence of oleic acid, whereas oleic acid supplementation induced filamentous growth by day 2 and maintained it through day 6 (24). The yeast-associated condition showed increased TGF-β and reduced IL-18 during day 2-6, consistent with a regulatory or resolution-associated response. Under filament-inducing conditions, both signals were reduced, suggesting a mixed or altered immune response (Supplementary Fig. S8, A).

A second study examined primary keratinocytes exposed to conditioned media from different *Malassezia* species (25). *M. globosa*-conditioned medium was associated with time-dependent reductions in IL12A, IL12B, IL10, and IL6 and increased IL18 and IL1A expression, whereas *M. restricta*-conditioned medium reduced IL12A and IL12B while increasing IL10 (Supplementary Fig. S8, B). These patterns are consistent with relatively inflammatory and regulatory response profiles, respectively. *M. globosa* has documented pseudohyphal-forming capacity, whereas comparable morphogenesis has not been reported for *M. restricta*.

A third study examined immune responses to heat-killed *M. pachydermatis*, which remains in the yeast form (26). Exposure to the yeast preparation did not strongly polarize macrophages, whereas subsequent stimulation elicited responses enriched in IL-10 and IL-4. This pattern is compatible with the resolution-permissive responsiveness associated with yeast-derived extracellular environments in our experiments.

Viewed together, these datasets are consistent with a state-dependent model of fungal-host interaction proposed in our study. Predominantly yeast-forming populations may favor regulatory or resolution-associated responsiveness; populations undergoing an unsynchronized yeast-to-pseudohyphal transition may generate mixed inflammatory and regulatory signals; and populations that establish a stable pseudohyphal state may bias host responses toward inflammation.

## Discussion

### Reframing fungal morphology

Fungal morphology is commonly interpreted through developmental and pathogenic frameworks, particularly those emphasizing the invasive properties of true hyphae (7–17). Our findings support an additional perspective: yeast and pseudohyphal forms can represent distinct physiological states whose metabolic organization influences the extracellular environment encountered by host cells. In this context, pseudohyphal growth should not be treated simply as an intermediate between yeast and true hyphae. Instead, it may reflect an adaptive state with direct consequences for how fungi interact with surrounding tissues and inflammatory signals.

### Pseudohyphal growth as an adaptive metabolic state

The temporal organization of the pseudohyphal response suggests a coordinated redistribution of metabolic priorities. Early changes in one-carbon metabolism, methyl-donor-associated pathways, and amino acid utilization were followed by remodeling of sulfur-associated metabolism and expansion of glutathione biosynthesis. Rather than representing isolated responses to culture stress, these changes are consistent with a transition linking nutrient utilization, redox management, and antioxidant defense.

Sustained expression of genes associated with mitochondrial adaptation, genome maintenance, and glutathione synthesis further suggests that pseudohyphal cells maintain a physiological program distinct from that of yeast cells. One interpretation is that these responses help accommodate altered redox demands when tryptophan-aldehyde metabolism is attenuated. However, the present data establish temporal association rather than a direct compensatory mechanism, and targeted perturbation studies will be needed to determine whether these pathways initiate, reinforce, or merely accompany the pseudohyphal state.

This distinction has practical implications for interpreting fungal stress responses. A stable pseudohyphal population may retain a coordinated metabolic configuration after the initiating environmental conditions have changed, allowing its extracellular effects to differ from those of a genetically identical yeast population exposed to the same subsequent environment. The persistence and reversibility of this configuration, and the mechanisms that maintain it, remain important questions for future investigation.

Changes in methionine and sulfur metabolism also raise the possibility that metabolic adaptation is linked to chromatin-associated regulation. Methionine metabolism determines the availability of SAM, the principal methyl donor for many cellular methylation reactions, and could therefore connect nutrient and redox status to histone modification or other chromatin-dependent processes. Our measurements do not demonstrate DNA methylation, histone methylation, or epigenetic stabilization. Nevertheless, testing whether altered methyl-group availability influences chromatin state may clarify how transient environmental pressures contribute to sustained pseudohyphal physiology.

### Tryptophan-aldehyde metabolism as a potential transition-associated node

Tryptophan-derived products such as IAA and TOL are often considered primarily as extracellular signaling or quorum-associated molecules (27, 28). Our findings suggest that their production also reflects an interconnected aldehyde-processing network with implications for intracellular redox balance. In particular, ALDH-mediated oxidation of IAAld to IAA links aldehyde clearance with NAD⁺ reduction and distinguishes the metabolically active yeast state from the pseudohyphal state. The attenuation of this branch could therefore mark, or potentially contribute to, a shift in metabolic organization during morphogenesis. Whether altered ALDH activity precedes the transition, results from it, or participates in a feedback loop remains unresolved. Defining the regulatory inputs controlling the ALDH-IAA axis will be essential for determining whether it functions as a causal checkpoint or as a marker of fungal physiological state.

### A state-dependent model of fungal-host interaction

The host-cell experiments indicate that morphotype-associated extracellular environments do not directly impose canonical inflammatory or tissue-repair macrophage polarization. Instead, they alter how innate immune cells respond to subsequent stimuli: yeast-associated environments favor responsiveness to tissue-repair cues, whereas pseudohypha-associated environments favor responsiveness to inflammatory cues. The inability of selected individual metabolites to reproduce the complete conditioned-media response further suggests that this activity depends on the combined extracellular milieu rather than a single measured compound. This conditioning model places fungal metabolism upstream of context-dependent immune responsiveness rather than equating fungal exposure with an immediate, fixed immune phenotype.

This framework may help explain why *Malassezia*-associated skin disease has been described as involving persistent inflammation in some settings and mixed inflammatory-regulatory responses in others (27–29). Predominantly yeast-forming populations may support commensal behavior and favor inflammatory resolution. During an unsynchronized yeast-to-pseudohyphal transition, the coexistence of yeast and pseudohyphal states could generate a composite extracellular environment that elicits both inflammatory and regulatory responses. If the pseudohyphal state becomes stabilized within a permissive host environment, sustained exposure to its associated extracellular products may instead bias local immune responsiveness toward persistent inflammation. Although this model requires further experimental validation, it offers a coherent interpretation of previously reported findings that have appeared inconsistent or difficult to reconcile (24–26, 29, 30).

#### Clinical implications and limitations

These observations suggest that morphogenetic potential, population synchrony, and metabolic state may provide information beyond fungal detection or abundance when evaluating host-fungal interactions. However, our conclusions are based on controlled fungal cultures, extracellular metabolite measurements, and a THP-1 model rather than clinical specimens. Noted that direct interpretation of intracellular metabolite pools is complicated in *Malassezia* by the difficulty of recovering intracellular material without cell-wall disruption or leakage, making extracellular trajectories an informative representation of cellular metabolism. Future studies should test defined clinical isolates in matched primary immune cells and skin models, establish the causal role of the ALDH-IAA pathway, and determine whether methionine-dependent chromatin regulation contributes to persistence of the pseudohyphal state.

#### Conceptual synthesis

Together, our findings connect fungal morphology, metabolic organization, and context-dependent innate immune responsiveness within a unified state-dependent framework. Yeast and pseudohyphal growth are associated with different extracellular environments that influence whether subsequent host signals are more likely to produce resolution-associated or inflammatory responses. Considering fungal functional state alongside host context may therefore improve interpretation of *Malassezia*-associated tissue homeostasis and chronic inflammatory disease.

## Materials and Methods

### Fungal strains and synchronized culture platform

*Malassezia furfur* strain CBS1878 (ATCC 14521) was used throughout this study. A single colony isolated from modified Leeming–Notman agar (MLNA) (31) was expanded in lipid-supplemented yeast extract-peptone-dextrose (YPD) medium at 32°C. The resulting yeast culture was divided into two populations. One population was maintained under yeast-growth conditions, whereas the second population was transferred into induction medium (IM) to generate pseudohyphal cells (14–17). Morphological states were verified by bright-field microscopy prior to downstream analyses. To compare metabolic adaptation from two pre-established fungal metabolic states, yeast and pseudohyphal cultures derived from the same clonal population were normalized to equivalent cell densities and transferred into identical serum-free RPMI medium. Culture supernatants were collected at defined time points for temporal metabolomic, isotope-tracing, and targeted metabolite analyses. Because both morphotypes originated from the same genetic background and were subsequently exposed to an identical nutrient environment, the resulting metabolic trajectories reflect intrinsic differences in metabolic organization associated with the pre-established fungal metabolic states rather than differences in genotype or extracellular nutrient availability. This synchronized culture platform enabled direct comparison of temporal metabolic adaptation between the yeast-associated and pseudohyphal-associated fungal metabolic states. The experimental workflow is summarized in Fig.1, B–D.

### Temporal metabolomic profiling

Cell-free culture supernatants were collected at 1, 8, and 24 h following transfer into RPMI medium, clarified through 0.45-μm filters, and stored at −80°C until analysis. Untargeted metabolomic profiling was performed using the Biocrates MxP® Quant 500 XL platform (32, 33). Quantitative metabolite concentrations were used for correlation-based trajectory analysis, pathway analyses, and metabolite ratio analyses. The complete metabolomic dataset is provided in Supplementary Figure S1, Table S1, and representative metabolite trajectories are presented in Supplementary Fig. S2.

### Stable isotope tracing of tryptophan metabolism and LC-MRM analysis of tryptophan-derived metabolites

To quantify metabolic flux through tryptophan metabolism, synchronized yeast and pseudohyphal cultures were first preconditioned in RPMI medium containing unlabeled L-tryptophan for 2 h before transfer into RPMI supplemented with uniformly labeled U-¹³C₁₁-L-tryptophan (5 mg/L; Cambridge Isotope Laboratories; Cat. No. CLM-4290). Cell-free culture supernatants were collected at 2, 15, 24, and 48 h, clarified by centrifugation (3,000 × g, 5 min) and filtration through 0.22-µm membrane filters, and stored at −80°C until analysis. Stable isotope tracing experiments were designed to evaluate temporal carbon flux through tryptophan-derived metabolic pathways using established principles of isotope-assisted metabolic flux analysis (34).

Tryptophan and its downstream metabolites, including indole-3-lactic acid (ILA), tryptophol (TOL), indole-3-acetic acid (IAA), indole-3-acetonitrile (IAN), indole-3-acetamide (IAM), and tryptamine (TAM), were quantified using a previously validated liquid chromatography–multiple reaction monitoring (LC–MRM) method with minor modifications for fungal culture supernatants (30). Briefly, samples were extracted by methanol precipitation in the presence of an isotopically labeled internal standard, dried under vacuum, reconstituted in 0.1% formic acid, and analyzed by LC–MRM. Chromatographic separation was performed on a Kinetex C18 column (2.1 × 100 mm, 2.6 µm; Phenomenex) coupled to an Agilent HPLC interfaced with a Sciex 3200 QTRAP mass spectrometer operated in positive electrospray ionization mode. Quantification was performed using authentic analytical standards and external calibration curves and uniformly labeled isotopologues were quantified using the corresponding unlabeled calibration curves as previously described (35). Isotopologue distributions were determined relative to unlabeled controls to evaluate temporal carbon flux through tryptophan-derived metabolic pathways.

### Stable isotope tracing and PRM-MS analysis of glutathione metabolism

To evaluate glutathione biosynthesis, synchronized yeast and pseudohyphal cultures were preconditioned in RPMI containing unlabeled glycine for 2 h before transfer into RPMI supplemented with uniformly labeled ¹³C₂-glycine (10 mg/L; Cambridge Isotope Laboratories, CLM-136). Cell-free culture supernatants were collected at 2, 15, 24, and 48 h, clarified through 0.45-μm filters, and stored at −80°C until analysis.

Reduced glutathione (GSH) and oxidized glutathione (GSSG) were quantified by parallel reaction monitoring mass spectrometry (PRM-MS) (36). Briefly, 10 μL of culture supernatant was derivatized with 100 mM iodoacetamide to convert GSH to carboxyamidomethyl-GSH, extracted with acetonitrile, evaporated to dryness, and reconstituted in 2% trifluoroacetic acid. Samples were analyzed using a Waters ACQUITY UPLC system coupled to a Thermo Scientific Q Exactive Plus mass spectrometer operated in positive electrospray ionization mode. Chromatographic separation was performed on an ACQUITY Premier HSS T3 column (50 × 2.1 mm, 1.8 μm) using a gradient of water and acetonitrile containing 0.1% formic acid. GSH and GSSG were quantified by PRM using compound-specific precursor and fragment-ion transitions with external calibration. Isotopically labeled glutathione species were quantified using the corresponding unlabeled calibration curves.

### RNA extraction and quantitative RT-PCR

Total RNA was isolated from synchronized yeast and pseudohyphal cultures collected at 2, 15, 48, and 72 h following transfer into RPMI medium using the Quick-RNA Fungal/Bacterial Miniprep Kit (Zymo Research; Cat. No. R2014). First-strand cDNA was synthesized using the High-Capacity cDNA Reverse Transcription Kit (Applied Biosystems; Cat. No. 4368814). Quantitative RT–PCR was performed using PowerUp SYBR Green Master Mix (Thermo Fisher Scientific; Cat. No. A25777) on a real-time PCR system. Relative transcript abundance was normalized to ACT1 and β-tubulin, and gene expression was calculated using the comparative ΔΔCt method (37). Primer sequences are listed in Supplementary Table S2.

### Preparation of fungal extracellular metabolic environments

To generate extracellular metabolic environments for host-cell experiments, cell-free conditioned media were collected from synchronized yeast and pseudohyphal cultures 36 h after transfer into RPMI medium, corresponding to the establishment of their respective fungal metabolic states. Culture supernatants were clarified through 0.45-μm filters, lyophilized, and stored at −80°C until use. Before THP-1 cell culture experiments, lyophilized samples were reconstituted in fresh RPMI medium to their original volume (CM1) or diluted 1:1 with fresh RPMI (CM2). Throughout this study, these preparations are referred to as fungal extracellular metabolic environments, reflecting the integrated extracellular metabolite compositions generated by the synchronized yeast- and pseudohyphal-associated fungal metabolic states.

To evaluate whether individual metabolites were sufficient to reproduce the biological activities of the fungal extracellular metabolic environments, indole-3-acetic acid (IAA) (Cat. No. I2886), indole-3-lactic acid (ILA) (Cat. No. I5508), tryptophol (TOL) (Cat. No. T90301), γ-Aminobutyric acid (GABA) (Cat. No. A2129) reduced glutathione (GSH) (Cat. No. G4251), and oxidized glutathione (GSSG) (Cat. No. G4376) were prepared individually or in the indicated combinations at concentrations corresponding to twofold (2×) the highest concentrations measured during temporal metabolomic profiling and used for THP-1 stimulation under identical culture conditions. Unless otherwise indicated, all these metabolites were purchased from Sigma-Aldrich.

### THP-1 cell culture and macrophage reference states

THP-1 human monocytic cells (ATCC TIB-202) were maintained in RPMI-1640 medium supplemented with 10% heat-inactivated fetal bovine serum (Thermo Fisher Scientific; Cat. No. A5256701), 1% penicillin–streptomycin (Cat. No. 15140122), and L-glutamine (Cat. No. 25030081) at 37°C in a humidified atmosphere containing 5% CO₂. Cytokine-defined reference populations were established using 20 ng/mL recombinant human GM-CSF (Cat. No. 300-03) or 25 ng/mL recombinant human M-CSF (Cat. No. 300-25) to generate macrophage maturation controls, followed by 20 ng/mL GM-CSF plus 100 ng/mL lipopolysaccharide (LPS) (Cat. No. 00-4976-93) to generate inflammatory reference states and 25 ng/mL M-CSF plus 20 ng/mL recombinant human IL-4 (Cat. No. 200-04) to generate tissue-repair reference states, as previously described (38–40). Unless otherwise indicated, all cytokines and LPS were purchased from Thermo Fisher Scientific. THP-1 cells were also cultured in yeast- or pseudohypha-derived extracellular metabolic environments before analysis of macrophage lineage-associated transcription factors, cytokine production, and cell-surface marker expression. For secondary stimulation experiments, THP-1 cells conditioned with yeast- or pseudohypha-derived extracellular metabolic environments were challenged with LPS (100 ng/mL), TNF-α (20 ng/mL), IL-4 (20 ng/mL), or TGF-β1 (20 ng/mL) for 6 h, after which culture supernatants were collected for cytokine analysis.

### Flow cytometry

THP-1 cells were washed with phosphate-buffered saline (PBS) containing 1% fetal bovine serum (FBS), stained with a Fixable Viability Dye (BD Pharmingen; Cat. No. 564406), and incubated with an Fc receptor blocking reagent before staining with fluorophore-conjugated antibodies against CD14 (BD Pharmingen; Cat. No. 567732), CD80 (Cat. No. 566992), CD86 (Cat. No. 560957), CD163 (Cat. No. 567882), and CD206 (Cat. No. 566884) according to the manufacturers’ recommended concentrations. Samples were analyzed using a BD LSRFortessa flow cytometer (BD Biosciences). Cell populations were first gated using forward scatter (FSC) and side scatter (SSC) to exclude debris, followed by doublet exclusion and viability gating before analysis of cell-surface marker expression. CD14 was used to evaluate progression from the monocyte state toward macrophage maturation, whereas CD80 and CD86 were analyzed as markers associated with inflammatory macrophage activation and CD163 and CD206 as markers associated with tissue-repair macrophage activation, consistent with current macrophage phenotyping guidelines. Data were analyzed using WinList 10 (Verity Software House, Topsham, ME, USA). Representative gating strategies are presented in Supplementary Figs. S5 and S6.

### Cytokine measurements

Culture supernatants were collected following the indicated stimulation protocols, and IL-10 (Thermo Fisher Scientific, Cat No. EHIL10) and IL-12p70 (Thermo Fisher Scientific, Cat No. BMS238) concentrations were quantified using human ELISA kits according to the manufacturer’s instructions. Cytokine production was used as a functional readout of the innate immune response state established by fungal extracellular metabolic environments following secondary stimulation with LPS, TNF-α, IL-4, or TGF-β.

### Statistical analysis

All experiments were performed using four independent biological replicates (n = 4) unless otherwise indicated. Data are presented as mean ± SD. Statistical analyses were performed using GraphPad Prism (version 10.1; GraphPad Software, Boston, MA, USA). Comparisons between two groups were analyzed using unpaired two-tailed Student’s t tests, whereas comparisons among multiple groups were analyzed using one-way analysis of variance (ANOVA) followed by the appropriate post hoc multiple-comparison test, as indicated in the corresponding figure legends. When a biological replicate met the predefined outlier criteria using the GraphPad Prism ROUT test (Q = 1%) or another predefined objective criterion, it was excluded from the analysis before statistical testing. Consequently, individual datasets contain either three or four biological replicates, as indicated by the plotted data points. Differences were considered statistically significant at P < 0.05.

## Data, Materials, and Software Availability

All data supporting the findings of this study are included in the article and its Supporting Information. Quantitative metabolomic data are provided in Supplementary Table S1, and primer sequences used for quantitative RT-PCR are provided in Supplementary Table S2. Additional data supporting the findings of this study are available from the corresponding author upon reasonable request.

## Acknowledgments

This work was supported by faculty startup funds awarded to Mei Zhang by Case Western Reserve University School of Medicine. We thank the Proteomics & Metabolomics Core at Case Western Reserve University School of Medicine and the Flow Cytometry Core at Case Comprehensive Cancer Center for their technical services and support.

## Author contributions

M.Z. conceived the study, secured funding, designed and supervised the research, interpreted the data, and wrote the manuscript. H.K. performed experiments, collected samples, analyzed data, contributed to the *Malassezia furfur* transition design, and contributed to the manuscript writing. M.N. performed experiments, collected samples, conducted analysis, contributed to the data interpretation, and contributed to manuscript writing. C.W. contributed to experimental design. S.V. contributed to the design and execution of the stable isotope-tracing experiments. M.M. contributed to the design of the tryptophan isotope-tracing experiments and the relevant data interpretation. S.K. contributed to the *Malassezia furfur* transition design and interpretation of pseudohyphal transition data. All authors reviewed, edited, and approved the final manuscript.

## Competing interests

The authors declare no competing interests.

---

**Supplementary Fig. S1.**
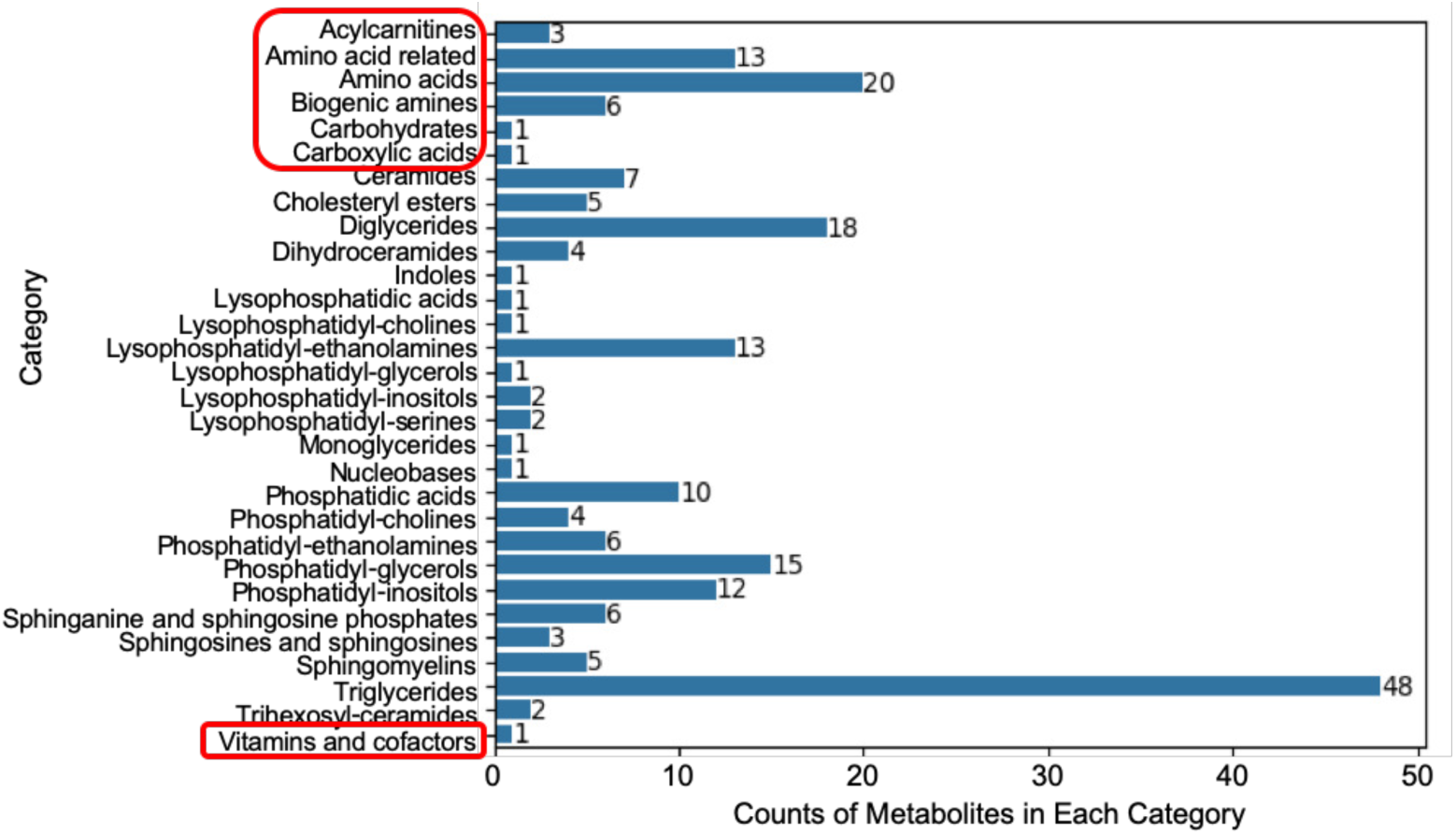
Overview of metabolite classes detected in temporal metabolomic profiling of *Malassezia furfur*. Metabolomic profiling was performed using the Biocrates MxP® Quant 500 XL platform. Bars indicate the number of metabolites detected within each metabolite class across yeast and pseudohyphal cultures. The metabolite classes highlighted by the red box represent the major categories investigated in this study. Individual metabolite identities, annotation information and concentrations are provided in Supplementary Table S1.

**Supplementary Fig. S2.**
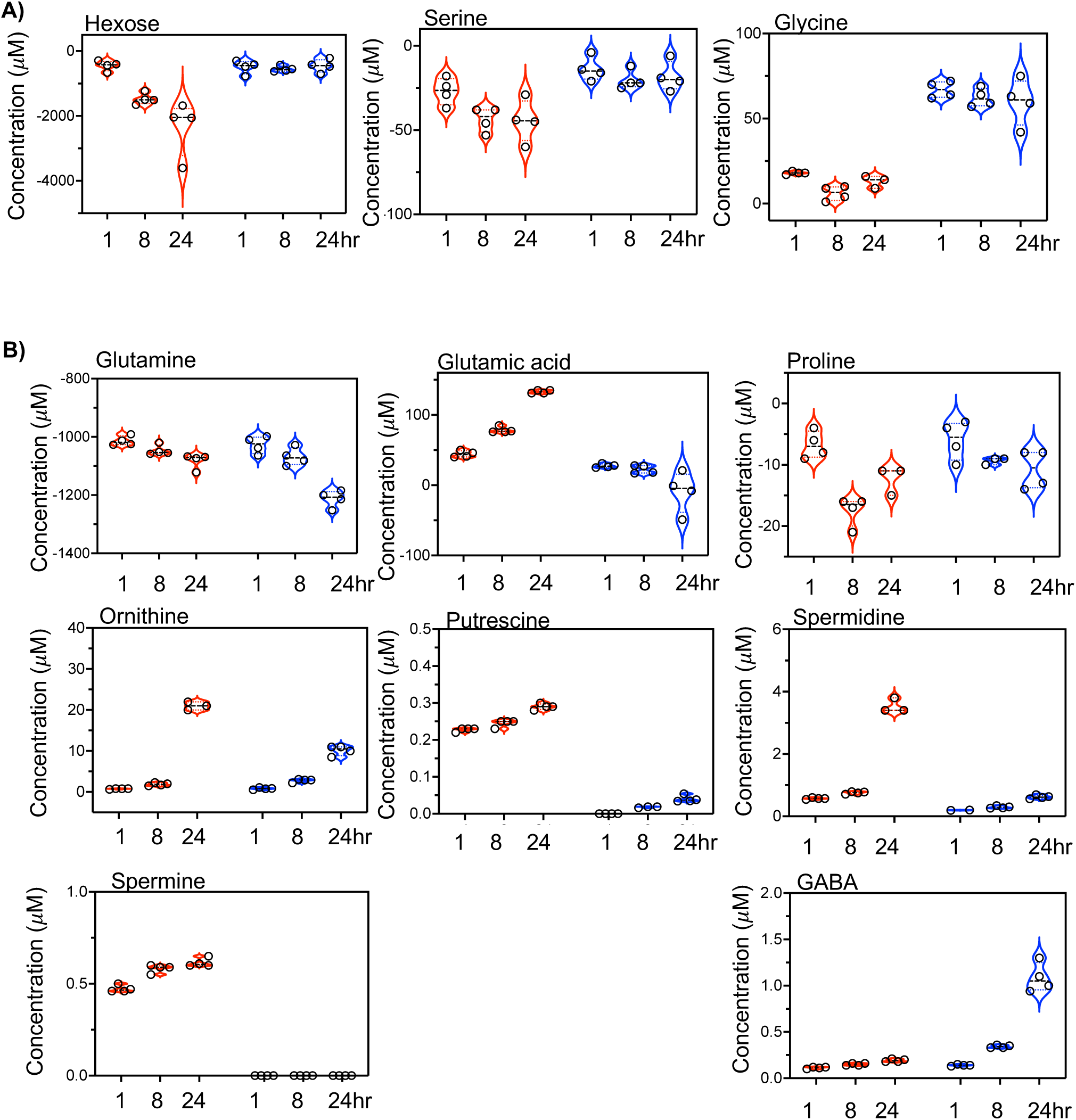
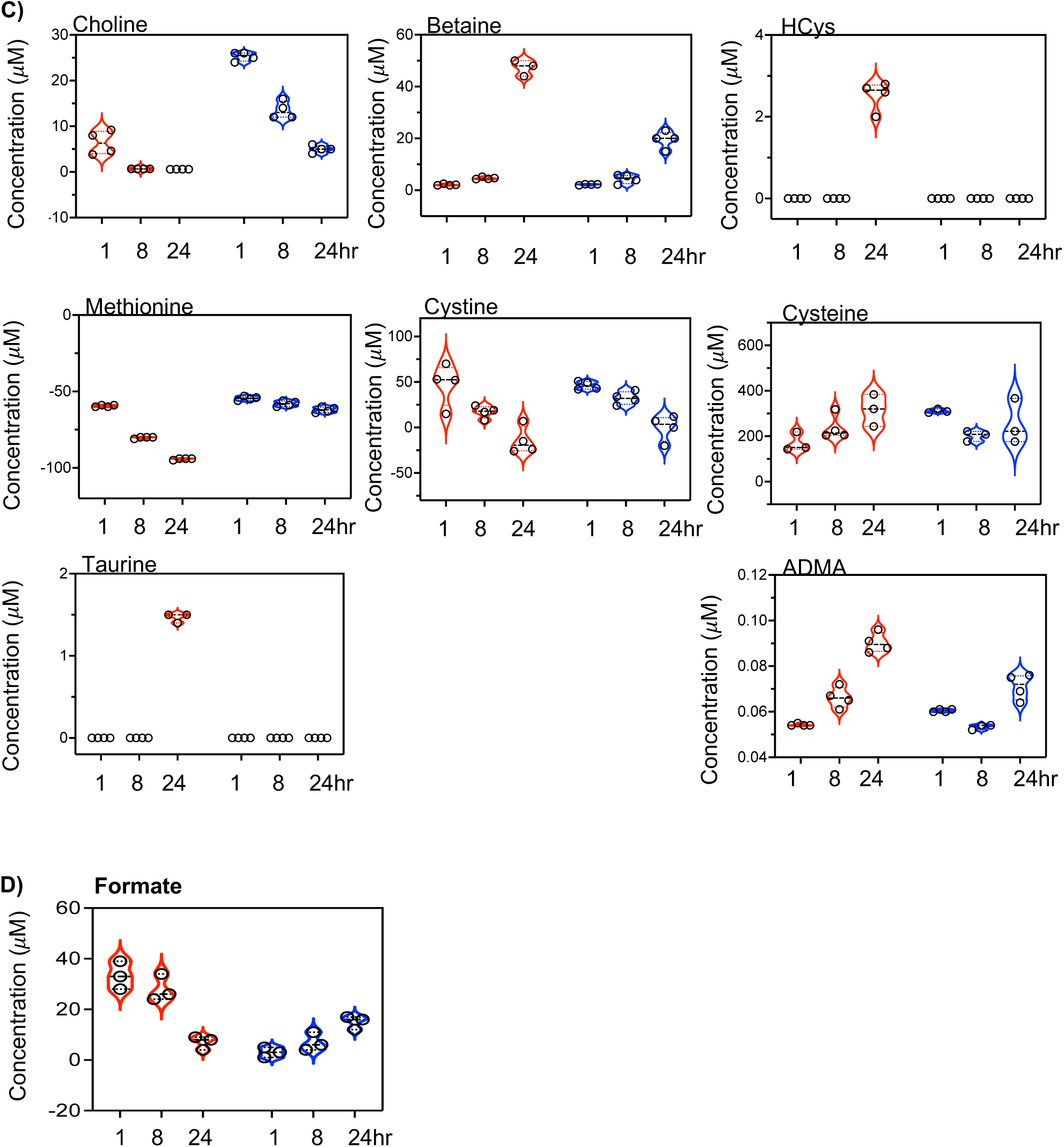
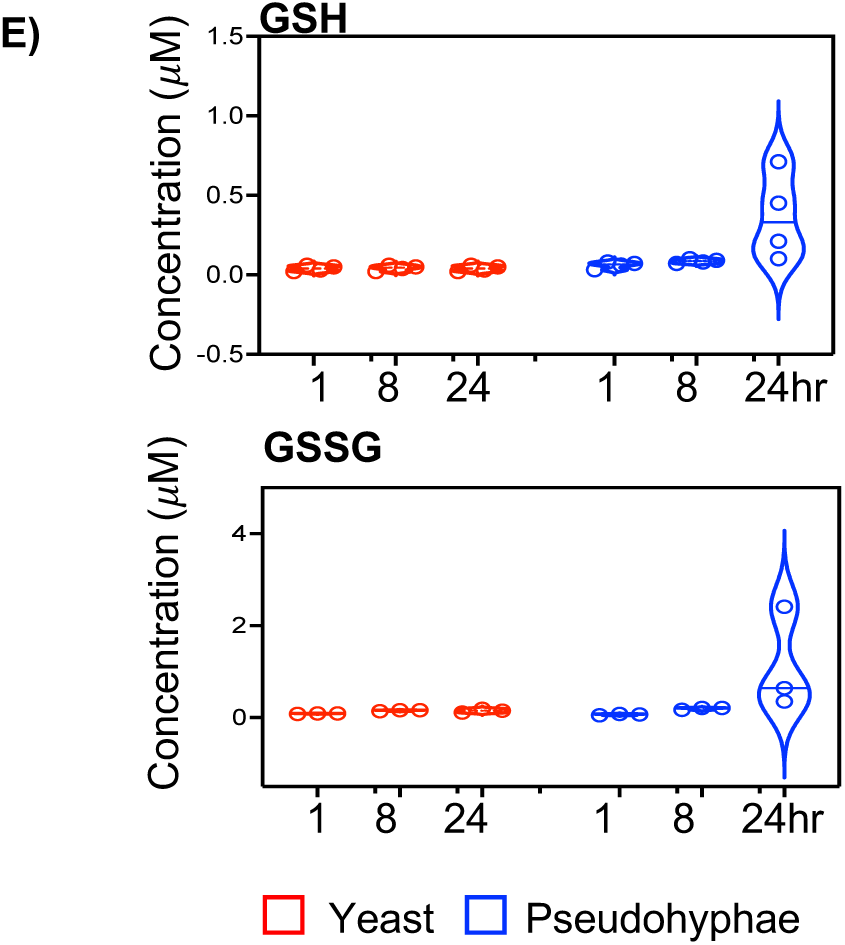
Representative temporal metabolite trajectories associated with yeast- and pseudohyphal-associated fungal metabolic states. **(A–C)** Representative metabolites identified by temporal metabolomic profiling of *M. furfur*. Metabolite abundances were quantified at 1, 8, and 24 h following transfer of established yeast and pseudohyphal cultures into serum-free RPMI. The selected metabolites illustrate temporal trajectories within major metabolic pathways identified by the global trajectory analysis in Fig. 1D, including one-carbon metabolism, amino acid metabolism, methionine cycling, sulfur amino acid metabolism, polyamine metabolism, and glutathione-associated pathways. **(D)** Targeted LC–MS/MS quantification of formate during nutrient utilization. **(E)** Targeted LC–MS/MS quantification of reduced glutathione (GSH) and oxidized glutathione (GSSG) during nutrient utilization.

## Supplementary Results

### Representative metabolic trajectories reveal coordinated remodeling of interconnected metabolic pathway

Metabolite trajectories were examined to determine how synchronized yeast and pseudohyphal cultures reorganized metabolism following transfer into a common nutrient environment (Supplementary Fig. S2).

Rather than displaying isolated changes in individual metabolites, multiple interconnected pathways underwent coordinated temporal remodeling, providing the experimental basis for the integrated pathway model presented in Fig. 1E.

Yeast cultures exhibited rapid utilization of glycine and serine together with progressive depletion of extracellular hexoses (Supplementary Fig. S2A,B). Coordinated changes in serine/glycine utilization, homocysteine–methionine cycling, taurine production, and polyamine metabolism—including the ornithine–putrescine–spermidine–spermine pathway—were consistent with remodeling of a folate-linked one-carbon network supporting S-adenosylmethionine (SAM)-dependent methyl-group metabolism (Supplementary Fig. S2A–C). Concurrent decreases in extracellular formate, independently quantified by targeted LC–MS/MS, further supported remodeling of one-carbon metabolism during nutrient adaptation (Supplementary Fig. S2D). Collectively, these temporal trajectories are consistent with establishment of a biosynthetically active metabolic organization during proliferative yeast growth.

In contrast, pseudohyphal cultures exhibited comparatively reduced serine/glycine utilization together with increased cysteine utilization and progressive expansion of reduced and oxidized glutathione pools (GSH and GSSG) (Supplementary Fig. S2A–E). Targeted LC–MS/MS independently confirmed increased GSH and GSSG abundance during later stages of nutrient utilization (Supplementary Fig. S2E). In parallel, increased extracellular γ-aminobutyric acid (GABA), together with comparatively attenuated polyamine-associated trajectories, suggested reorganization of nitrogen-associated metabolism. These representative trajectories are therefore consistent with coordinated remodeling toward glutathione-associated antioxidant metabolism and redox homeostasis during stabilization of the pseudohyphal morphotype.

Collectively, the representative metabolite trajectories demonstrate coordinated remodeling across one-carbon metabolism, methionine cycling, transmethylation reactions, polyamine metabolism, sulfur amino acid metabolism, glutathione metabolism, and central carbon metabolism. These analyses provide the experimental basis for the integrated pathway model presented in Fig. 1E.

**Supplementary Fig. S3.**
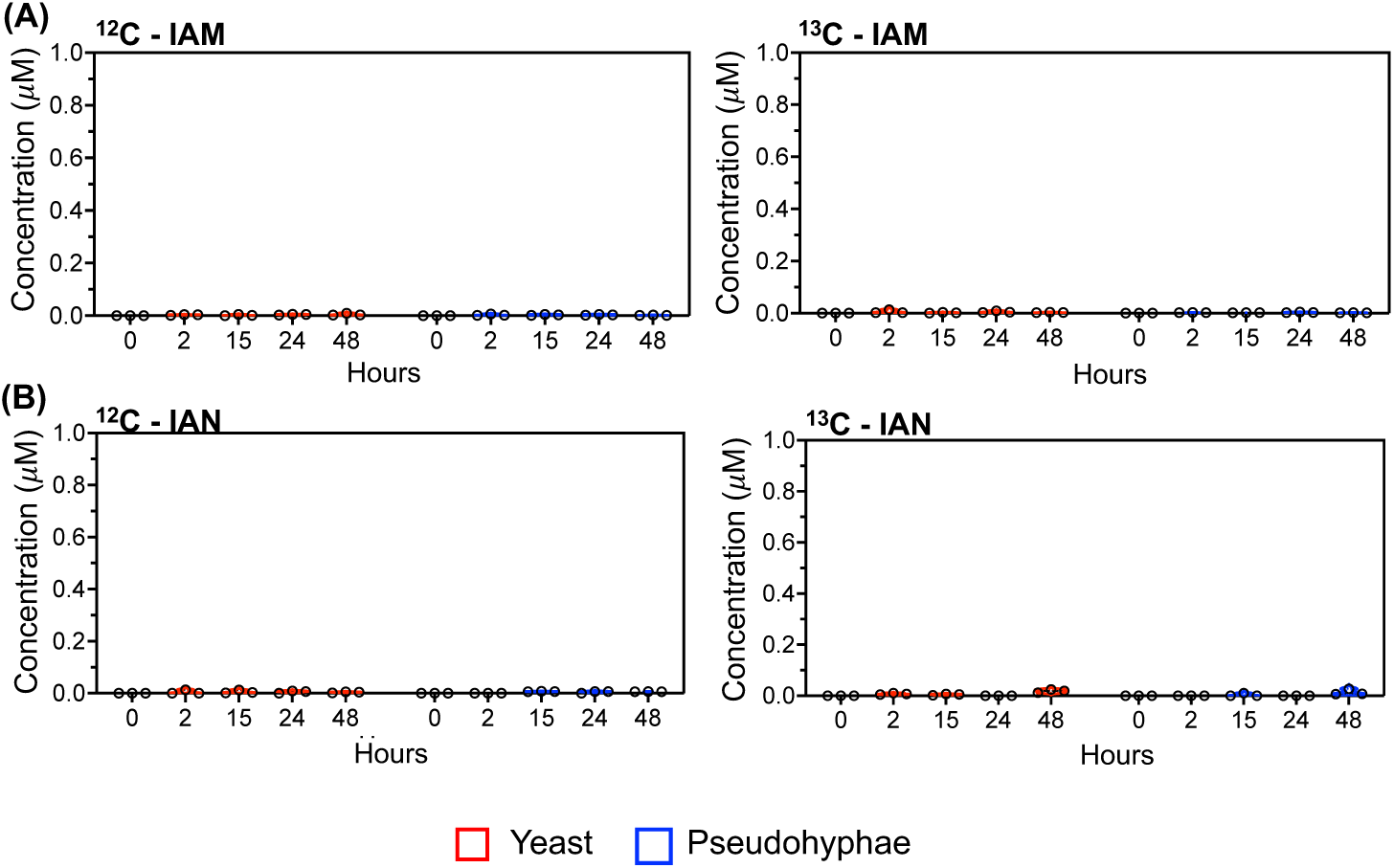
Stable isotope tracing demonstrates the absence of detectable indole-3-acetamide and indole-3-acetonitrile pathway activity during tryptophan metabolism in *M. furfur*. Established yeast and pseudohyphal cultures were transferred into serum-free RPMI supplemented with uniformly labeled ¹³C₁₁-tryptophan. Endogenous (¹²C) and newly synthesized (¹³C) metabolites were quantified by LC–MS/MS at the indicated time points. **(A)** Quantification of indole-3-acetamide (IAM) and related metabolites. **(B)** Quantification of indole-3-acetonitrile (IAN) and related metabolites. Neither endogenous nor ¹³C-labeled IAM or IAN was detected in extracellular space under the experimental conditions. These findings support the pathway organization illustrated in Fig. 3B.

**Supplementary Fig. S4.**
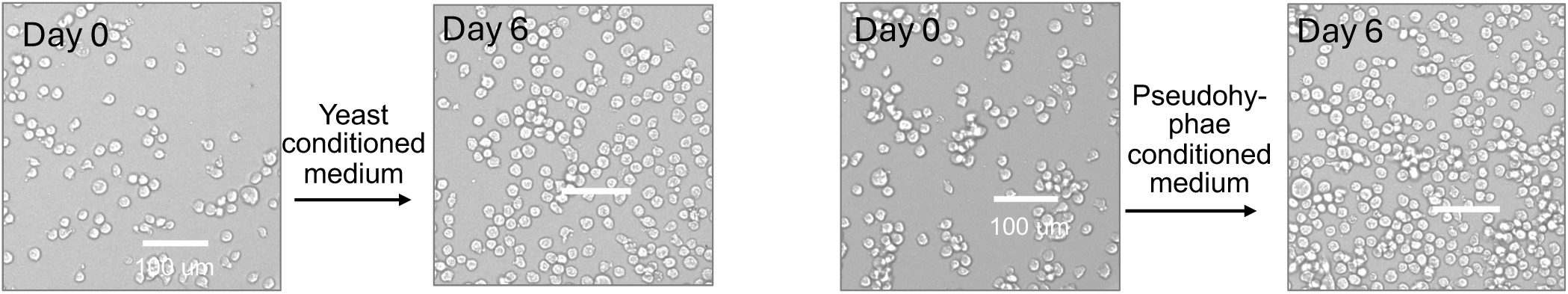
Morphological assessment of THP-1 cells cultured with fungal-conditioned media. Representative bright-field images of THP-1 cells cultured for six days in RPMI alone or in extracellular metabolic environments generated by established yeast or pseudohyphal *M. furfur* cultures. No overt morphological evidence of cytotoxicity or canonical macrophage differentiation was observed following exposure to either extracellular metabolic environment.

**Supplementary Fig. S5.**
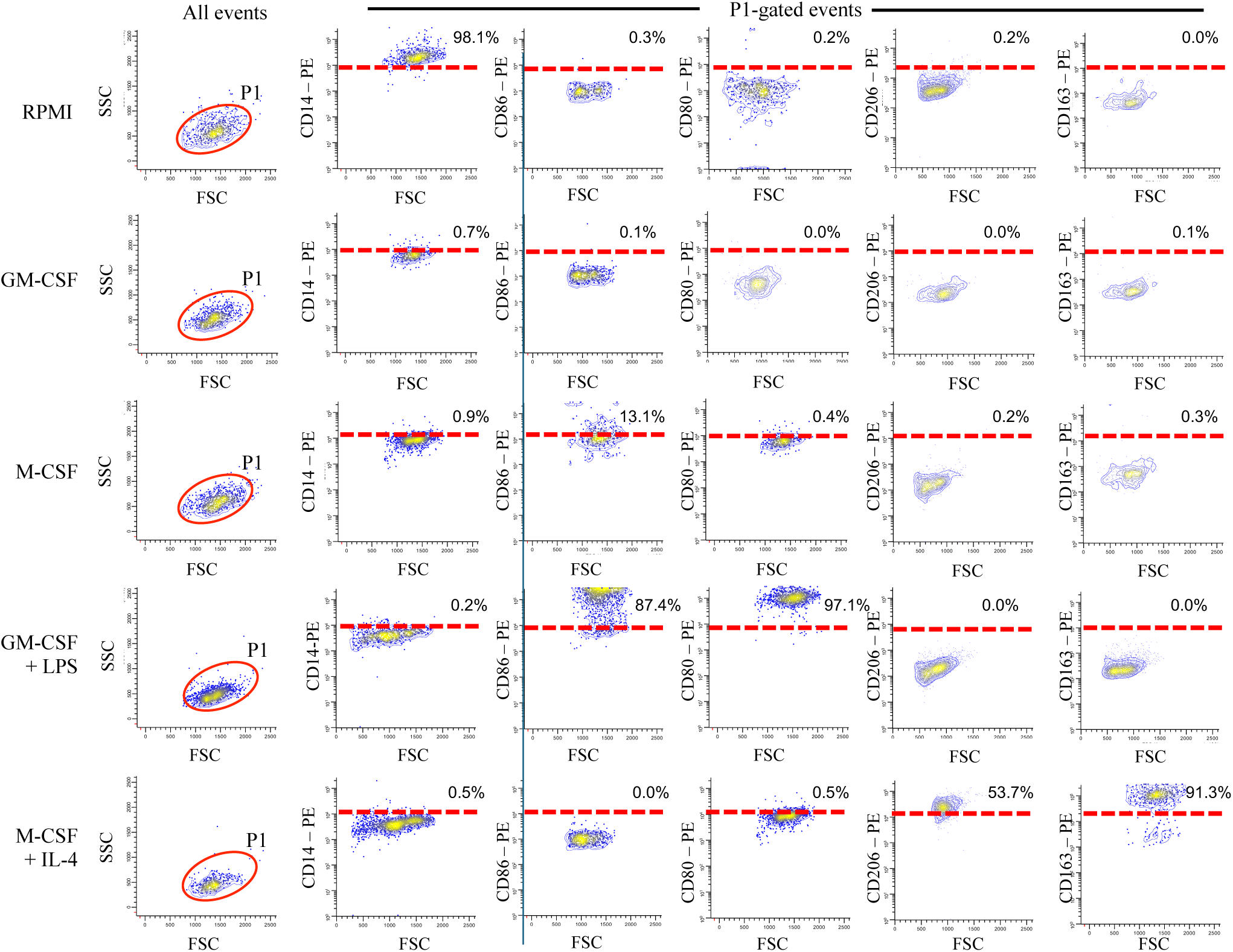
Flow cytometric characterization of THP-1 maturation and polarization reference states in Fig. 5B. THP-1 cells were cultured under the indicated cytokine conditions to establish reference states for macrophage maturation and activation. Experimental groups included RPMI alone (undifferentiated control), GM-CSF, M-CSF, GM-CSF plus LPS (pro-inflammatory reference), and M-CSF plus IL-4 (tissue-repair reference). Cells were first gated on the P1 population using forward scatter (FSC) and side scatter (SSC), followed by analysis of CD14, CD80, CD86, CD206, and CD163 expression. Red dashed lines indicate the positivity threshold for each marker. Representative flow cytometry contour plots from three independent experiments are shown. CD14 was evaluated as a marker of monocyte maturation, whereas CD80 and CD86 were analyzed as markers associated with inflammatory macrophage activation and CD206 and CD163 as markers associated with tissue-repair macrophage activation. These cytokine-defined reference populations were used for comparison with THP-1 cells cultured in yeast- and pseudohyphal-derived extracellular metabolic environments.

**Supplementary Fig. S6.**
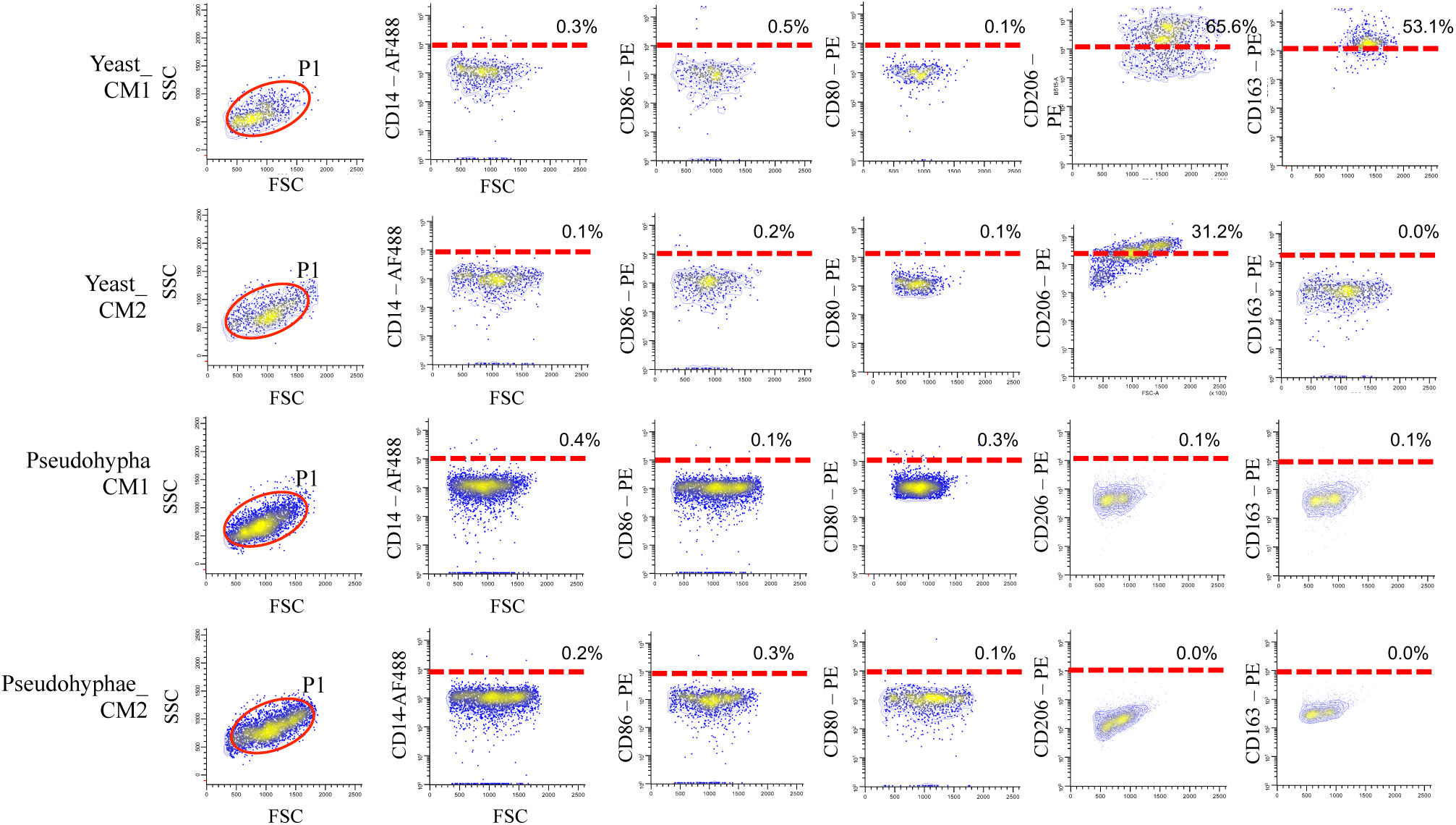
Flow cytometric characterization of THP-1 cells cultured in extracellular metabolic environments generated by *Malassezia furfur*. THP-1 cells were cultured in extracellular metabolic environments generated by established yeast and pseudohyphal *M. furfur* cultures under the same experimental conditions used for the lineage-associated gene expression and cytokine analyses presented in Fig. 5C. Extracellular metabolic environments were used either directly (CM1) or diluted 1:1 with fresh RPMI (CM2). Cells were first gated on the P1 population using forward scatter (FSC) and side scatter (SSC), followed by analysis of CD14, CD80, CD86, CD206, and CD163 expression. Red dashed lines indicate the positivity threshold for each marker. Representative flow cytometry contour plots from three independent experiments are shown. Marker definitions are identical to those described in Supplementary Fig. S4. These flow cytometric analyses complement the lineage-associated transcription factor (MAFB and SPI1) expression and cytokine measurements presented in Fig. 5C.

**Supplementary Fig. S7.**
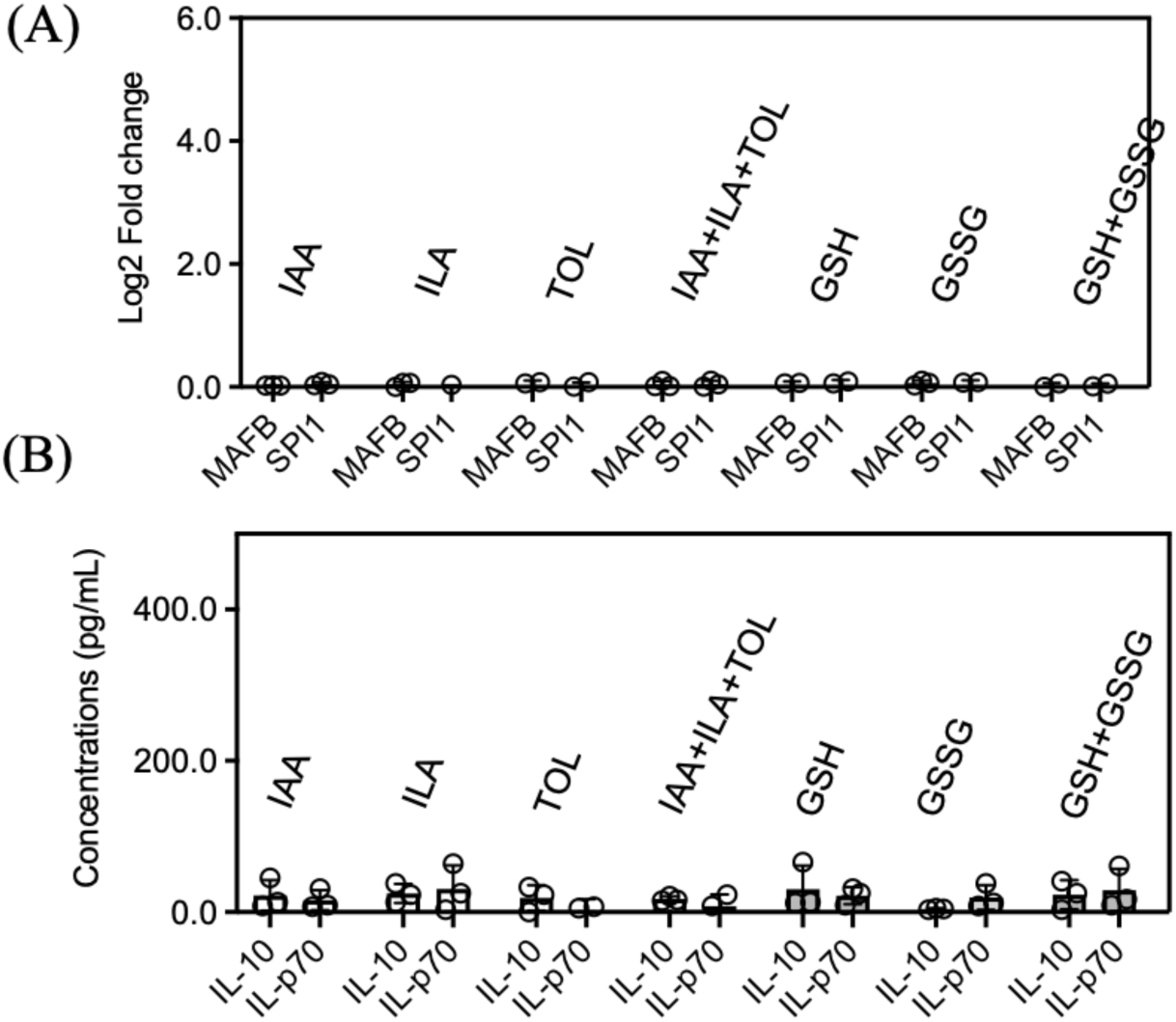
Individual metabolites and representative metabolite combinations do not reproduce the THP-1 maturation phenotype induced by *Malassezia furfur* extracellular metabolic environments. THP-1 cells were cultured with representative metabolites associated with the yeast- and pseudohyphal-associated metabolic programs of *M. furfur*, including IAA, ILA, TOL, GABA, GSH, and GSSG, either individually or in representative metabolite combinations, under the same experimental conditions used in Fig. 5D. Individual metabolites and metabolite combinations were tested at concentrations corresponding to twofold (2×) the highest metabolite levels measured by temporal metabolomic profiling. Macrophage lineage-associated transcription factor expression (MAFB and SPI1) and cytokine production (IL-10 and IL-12p70) were quantified by RT-qPCR and ELISA, respectively. Data represent three independent biological experiments (n = 3). None of the individual metabolites or tested metabolite combinations reproduced the transcriptional or cytokine responses induced by extracellular metabolic environments generated by yeast or pseudohyphal cultures.

**Supplementary Fig. S8.**
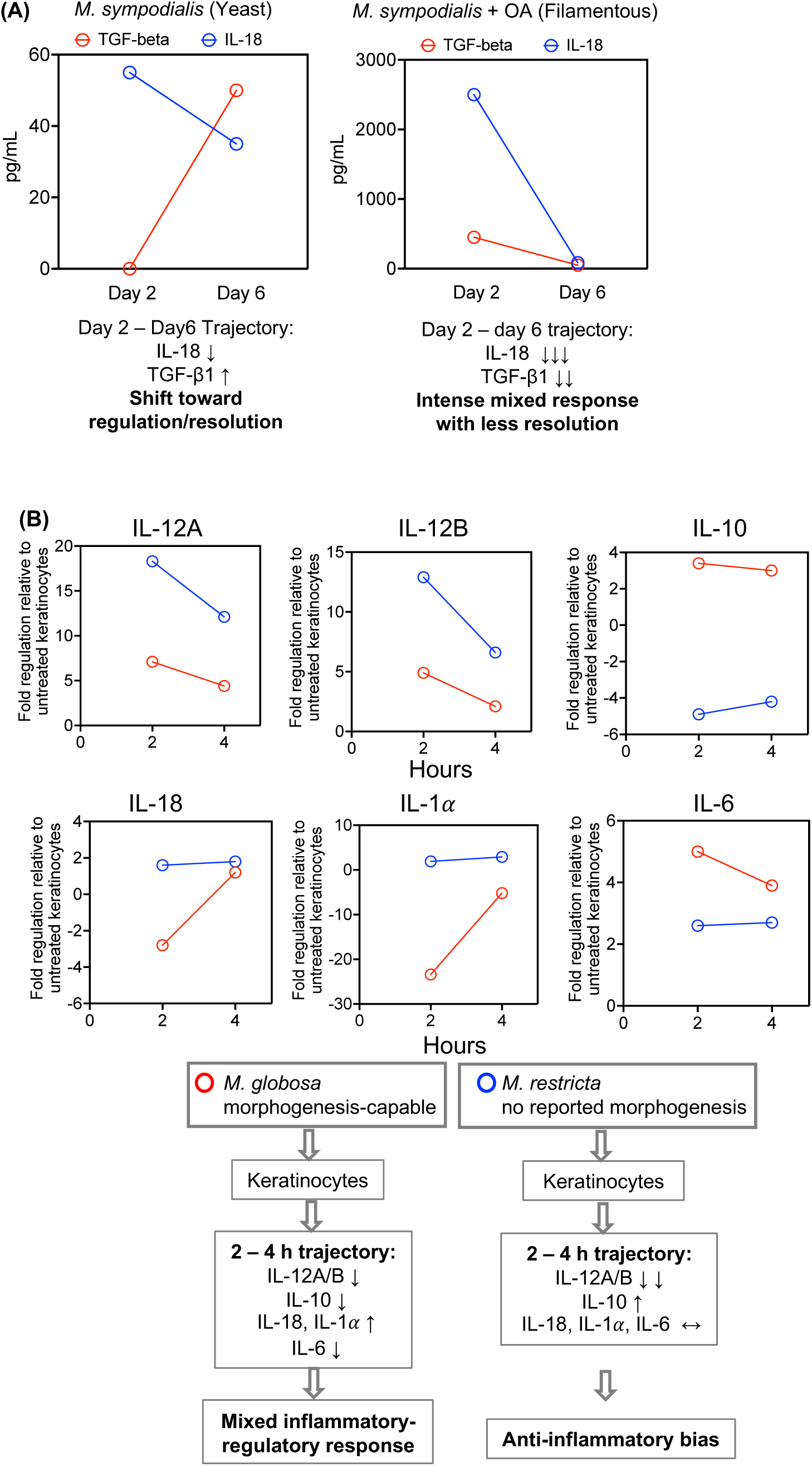
Independent trajectory reanalysis of published datasets supports a state-dependent model of *Malassezia*-host immune interactions. (A) Reanalysis of a human skin explant study (24) examining responses to *Malassezia sympodialis* under yeast- and filament-promoting conditions. In the absence of oleic acid, *M. sympodialis* remained predominantly in the yeast form and was associated with increased TGF-β and reduced IL-18 expression. Oleic acid supplementation induced filamentous growth and was associated with reduced TGF-β and IL-18 responses. (B) Reanalysis of keratinocyte responses to conditioned media from different *Malassezia* species (25). *M. globosa*-conditioned medium was associated with decreased IL12A, IL12B, IL10, and IL6 expression and increased IL18 and IL1A expression over time, whereas M. restricta-conditioned medium reduced IL12A and IL12B while increasing IL10 expression.

## References

1. Gaitanis G, Magiatis P, Hantschke M, Bassukas ID, Velegraki A (2012) Malassezia yeasts and their role in skin disease. J Eur Acad Dermatol Venereol 26:16–26.

2. Ashbee HR (2007) Update on the genus Malassezia. Med Mycol 45:287–303.

3. Saunders CW, Scheynius A, Heitman J (2012) Malassezia species on the skin: Commensals and pathogens. J Invest Dermatol 132:1465–1472.

4. Findley K, Oh J, Yang J, Conlan S, Deming C, Meyer JA, Schoenfeld D, Nomicos E, Park M, Program NCS, et al. (2013) Topographic diversity of fungal and bacterial communities in human skin. Nature 498:367–370.

5. Grice EA, Segre JA (2011) The skin microbiome. Nat Rev Microbiol 9:244–253.

6. Oh J, Byrd AL, Deming C, Conlan S, Kong HH, Segre JA (2014) Biogeography and individuality shape function in the human skin metagenome. Nature 514:59–64.

7. Sudbery PE (2011) Growth of Candida albicans hyphae. Nat Rev Microbiol 9:737–748.

8. Gimeno CJ, Ljungdahl PO, Styles CA, Fink GR (1992) Unipolar cell divisions in the yeast S. cerevisiae lead to filamentous growth. Cell 68:1077–1090.

9. Berman J (2006) Morphogenesis and cell cycle progression in Candida albicans. Curr Opin Microbiol 9:595–601.

10. Gow NAR, van de Veerdonk FL, Brown AJP, Netea MG (2012) Candida albicans morphogenesis and host defence: Discriminating invasion from colonization. Nat Rev Microbiol 10:112–122.

11. Noble SM, Gianetti BA, Witchley JN (2017) Candida albicans cell-type switching and functional plasticity in the mammalian host. Nat Rev Microbiol 15:96–108.

12. Youngchim S, Nosanchuk JD, Chongkae S, Vanittanokom N (2017) Ketoconazole inhibits Malassezia furfur morphogenesis in vitro under filamentation-optimized conditions. Arch Dermatol Res 309:47–53.

13. Dorn M, Roehnert K (1977) Dimorphism of Pityrosporum orbiculare in a defined culture medium. J Invest Dermatol 69:244–248.

14. Saadatzadeh MR, Ashbee HR, Holland KT, Ingham E (2001) Production of the mycelial phase of Malassezia in vitro. Med Mycol 39:487–493.

15. Barchmann T, Hort W, Krämer HJ, Mayser P (2011) Glycine as a regulator of tryptophan-dependent pigment synthesis in Malassezia furfur. Mycoses 54:17–22.

16. Mayser P, Tows A, Krämer HJ, Weiß R, Gaitzsch J (1998) Further characterization of Malassezia furfur morphogenesis and nitrogen-dependent filament formation. Mycoses 41:281–285.

17. Mayser P, Imkampe A, Winkeler M, Papavassilis C (1998) Growth requirements and nitrogen metabolism of Malassezia furfur. Arch Dermatol Res 290:277–282.

18. Chen H, Fink GR (2006) Feedback control of morphogenesis in Candida albicans by aromatic alcohols. Genes Dev 20:1150–1161.

19. Alonso-Roman R, Last A, Mirhakkak MH, et al. (2022) Lactobacillus rhamnosus colonisation antagonizes Candida albicans by forcing metabolic adaptations that compromise pathogenicity. Nat Commun 13:3192.

20. Pellon A, Begum N, Sadeghi Nasab SD, Harzandi A, Shoaie S, Moyes DL (2022) Role of cellular metabolism during Candida-host interactions. Pathogens 11:184.

21. Ene IV, Brunke S, Brown AJP, Hube B (2014) Metabolism in fungal pathogenesis. Cold Spring Harb Perspect Med 4:a019695.

22. Brown AJP, Brown GD, Netea MG, Gow NAR (2014) Metabolism impacts upon Candida immunogenicity and pathogenicity at multiple levels. Trends Microbiol 22:614–622.

23. Levy M, Thaiss CA, Elinav E (2016) Metabolites: Messengers between the microbiota and the immune system. Genes Dev 30:1589–1597.

24. Corzo-León DE, MacCallum DM, Munro CA (2021) Host responses in an ex vivo human skin model challenged with Malassezia sympodialis. Front Cell Infect Microbiol 10:561382.

25. Thomas DS, Ingham E, Bojar RA, Holland KT (2008) In vitro modulation of human keratinocyte pro- and anti-inflammatory cytokine production by the capsule of Malassezia species. FEMS Immunol Med Microbiol 54:203–214.

26. Ishibashi Y, Sugita T, Nishikawa A (2006) Cytokine secretion profile of human keratinocytes exposed to Malassezia yeasts. FEMS Immunol Med Microbiol 48:400–409.

27. Han TL, Cannon RD, Villas-Bôas SG (2011) The metabolic basis of Candida albicans morphogenesis and quorum sensing. Fungal Genet Biol 48:747–763.

28. Gushiken-Ibañez E, Stokmaier M, Barone G, Staropoli A, Karakaya T, Beer HD, Vinale F, Ianiri G, LeibundGut-Landmann S (2025) Commensal yeast Malassezia produces tryptophan metabolites to promote tissue homeostasis via the aryl hydrocarbon receptor in mice. Cell Host Microbe.

29. Saunte DML, Gaitanis G, Hay RJ (2020) Malassezia-associated skin diseases, the use of diagnostics and treatment. Front Cell Infect Microbiol 10:112.

30. Thayikkannu AB, Kindo AJ, Veeraraghavan M (2015) Malassezia: Can it be ignored? Indian J Dermatol 60:332–339.

31. Leeming JP, Notman FH (1987) Improved methods for isolation and enumeration of Malassezia furfur from human skin. J Clin Microbiol 25:2017–2019.

32. Wishart DS, Feunang YD, Marcu A, et al. (2018) HMDB 4.0: The Human Metabolome Database for 2018. Nucleic Acids Res 46:D608–D617.

33. Siskos AP, Jain P, Römisch-Margl W, et al. (2017) Interlaboratory reproducibility of a targeted metabolomics platform for analysis of human serum and plasma. Anal Chem 89:656–665.

34. Buescher JM, Antoniewicz MR, Boros LG, et al. (2015) A roadmap for interpreting 13C metabolite labeling patterns from cells. Nat Rev Mol Cell Biol 16:559–578.

35. Miyagi M, Uchihashi K, Hasegawa K, Akimoto T (2008) Simultaneous measurement of tryptophan and related compounds by liquid chromatography/electrospray ionization tandem mass spectrometry. J Chromatogr B Analyt Technol Biomed Life Sci 867:57–63.

36. Mathews TP (2023) Quantitation of glutathione and oxidized glutathione ratios from biological matrices using LC-MS/MS. Methods Mol Biol 2675:133–148.

37. Livak KJ, Schmittgen TD (2001) Analysis of relative gene expression data using real-time quantitative PCR and the 2−ΔΔCT method. Methods 25:402–408.

38. Murray PJ, Allen JE, Biswas SK, et al. (2014) Macrophage activation and polarization: Nomenclature and experimental guidelines. Immunity 41:14–20.

39. Fleetwood AJ, Lawrence T, Hamilton JA, Cook AD (2007) GM-CSF- and M-CSF-dependent macrophage phenotypes display differential dependence on transcription factor networks. J Immunol 178:5245–5252.

40. Martinez FO, Gordon S, Locati M, Mantovani A (2006) Transcriptional profiling of the human monocyte-to-macrophage differentiation and polarization: New molecules and patterns of gene expression. J Immunol 177:7303–7311.

